# ProxyTyper: Generation of Proxy Panels for Privacy-aware Outsourcing of Genotype Imputation

**DOI:** 10.1101/2023.10.01.560384

**Authors:** Degui Zhi, Xiaoqian Jiang, Arif Harmanci

## Abstract

One of the major challenges in genomic data sharing is protecting the privacy of participants. Numerous studies demonstrated that genetic data and related summary statistics can be used for identifying individuals. These studies led to a strong chilling effect on researchers that hindered collaborative data sharing. Consequently, population-level genetic databases are often siloed in central repositories with complex and burdensome data usage agreements. While cryptographic methods that are provably secure have been developed, these methods require high-level expertise in security and depend on large computational resources.

To fill the methodological gap in this domain, we present ProxyTyper, a set of data protection mechanisms to generate “proxy-panels” from sensitive haplotype-level genetic datasets. ProxyTyper uses haplotype sampling, allele hashing, and anonymization to protect the genetic variant coordinates, genetic maps, and chromosome-wide haplotypes. These mechanisms can provide strong deterrence against honest-but-curious entities and well-known re-identification and linking attacks. The proxy panels can be used as input to existing tools without any modifications to the underlying algorithms. We focus on building proxy-panels for genotype imputation to protect typed and untyped variants. We demonstrate that proxy-based imputation provides protection against well-known attacks with a minor decrease of imputation accuracy for variants in wide range of allele frequencies.

## Introduction

Decreasing cost of whole genome sequencing and genotyping brought about a massive increase in the number of personal genomes[1]. Starting with the initial population-scale and biobank sequencing[2] projects such as The HapMap Consortium[3,4], The 1000 Genomes Project[5] and the population scale projects such as UK Biobank, Genomics England, Trans-omics for precision medicine (TOPMed)[6], and *AllofUs* research program[7], millions of personal genomes are available. Local sequencing efforts from underrepresented populations which are vital for increasing diversity in research[8–10] and for inclusion of underrepresented populations[11,12]. These data provide resources for a large spectrum of genetic variants and create opportunities for understanding ancestral composition and mapping genetic risk factors of diseases by association studies.

However, sharing the genetic data brought up ethical challenges around privacy[13], e.g., usage for forensic purposes[14–16], and concerns about discrimination[17,18]. Public is becoming increasingly aware of the ethical and privacy implications around sharing genetic data, and regard privacy as one of the top factors while deciding who they share genetic data[19,20]. Numerous negative experiences (Havasupai Tribe[21,22], Ashkenazi Jews[23]) undermined the trust in the research community due to lack of ethical reporting standards and insensitivity to cultural norms. Due to these concerns, datasets are often siloed in the central server and are protected by data usage agreements and severely hamper collaborative research opportunities[24,25] even within the same consortia[26,27]. Although legislation such as GDPR in the EU[28] and HIPAA[29] in the USA aim at protecting personal data, their interpretation is not clear about genetic data. For example, GDPR requires using new technological methods to make sure that the data is “irreversibly de-identified” unless consent is available for sharing identifiable data. However, even when consent is granted for sharing genomes, individuals may not consent to the downstream characterization of their genomes[30].

The main concerns around genomic privacy[31–34] are related to the re-identification of an individual by “singling-out” (GDPR) within a cohort. Due to the high dimensional and complex correlative structure of genetic data [30,35,36], many routes are open for re-identification of individuals[33]. For example, only 100 carefully selected SNPs are sufficient to identify any individual[37]. The most famous attacks, including Homer’s t-statistics[38], Sankararaman’s LRT[39] demonstrated that an individual’s participation in a genome-wide association studies (GWAS) study can be reliably identified even when only allele frequencies of the study are known. These attacks were alarming at the time because of their simplicity and their applicability to summary statistics[40,41]. Further studies identified that participation in different studies can lead to linking attacks and reveal sensitive information[42,43] of participants and relatives[44,45]. Other attacks that make use of haplotype frequencies have been proposed to identify participation[46,47]. Recent studies demonstrated that genetic beacons[48] (where the existence of variants in a database are queried as yes/no answers) are vulnerable to Bustamante’s attacks[49] and more advanced attacks[50,51] where an adversary can identify participation of an individual in a beacon by adversarial querying, which may impact biobank scale databases. For example, the knowledge of the rare variant positions can be used to reveal participation in TOPMed panel using results obtained from TOPMed imputation server. One of the important factors that make these attacks possible is that they rely on a naive data-sharing model where raw data is made available to entities without a specific focus on a task such as genotype imputation. Furthermore, several studies showcased that the most well-known attacks are applied in very well-controlled scenarios and they lack of formal treatment of the false positive rates [52] when the assumptions are not satisfied [53].

By far, the most popular genetic data sharing model is the restricted access model that model relies on users signing agreements for each new dataset. Differential privacy[54,55] (DP) approaches design privacy-preserving data release mechanisms where aggregate statistics from data are computed and noise is added to the released statistics to enable privacy. While DP provides strong privacy guarantees, the noise addition is a major drawback since a naive noise addition may corrupt certain data types (such as genotype data) to the extent that the data become unusable. Encryption-based approaches are the most rigorous route to securely share genetic data albeit with low practicality and are applied to the field of genotype imputation[56,57], and association studies [58–61], database sequence queries [62], and read mapping[63,64]. Methods such as homomorphic encryption[65,66] enable analyzing and processing encrypted data directly. There are, however, a substantial computational (requirement for reformulation of algorithms under cryptographic primitives [67]), storage (ciphertext expansion), and maintenance costs. Moreover, each algorithm must be re-implemented and optimized to efficiently analyze the encrypted data. Multiparty computation (MPC) based systems [68,69] provide cryptographic security by dividing data among multiple parties. MPC-based systems rely on heavy communication and computational requirements on the parties [60] and require algorithms to be redesigned with careful reformulation and parameter selection (circuit depth, ciphertext size) to ensure that algorithms can be executed efficiently. Consequently, HE and MPC methods are not widely adopted in the community, yet. Trusted execution environments (Intel SGX, AMD SEV) use security-by-engineering which are vulnerable to numerous attacks[70,71] and rely on trust on a corporate entity. Of note, Intel SGX is currently deprecated for client-side processors. Although several anonymization techniques exist[72,73], they require reformulation of specific tasks (e.g., relative searching) and may not be easily applicable to different high throughput tasks.

Synthetic datasets have been explored as a potential approach that can be useful for protecting privacy[74] in collaborative studies. In these approaches, synthetic datasets are generated at each collaborating entity. In this framework, the entities generate a representative synthetic dataset using their local sensitive datasets. In the analysis, synthetic datasets are used instead of the original sensitive datasets. The privacy of participants is protected since the sensitive datasets are never used in analysis. In genomics, synthetic datasets are used for protecting reference panels in genotype imputation[75,76], and for analysis of ancestral simulations[77,78]. While these approaches provide means for generating synthetic datasets, privacy is still not explicitly introduced into the synthetic data generation models. For example, allele frequencies and coordinates are preserved in the synthetic data, which may make the synthetic datasets vulnerable to re-identification attacks.

Here, we present ProxyTyper, a framework for building “proxy panels”, i.e. panels that are similar in statistical properties to the original panel but are anonymized in terms of subjects, alleles, and variant coordinates. ProxyTyper utilizes three mechanisms to protect haplotype datasets in terms of variant positions, genetic maps, and variant genotypes: First mechanism protects the variant positions and genetic maps that can leak side-channel information. Second is resampling of original haplotype panels using a Li-Stephens Markov model with privacy parameters for tuning privacy level and utility. This step generates a mosaic of the original haplotypes so that each chromosome-wide haplotype is a mosaic of the haplotypes in the original panel. The third mechanism consists of encoding the alleles in resampled panels using locality-based hashing and permutation. One of the important aspects of proxy panels is that they can be used as input to existing tools. The proxy panels can be tuned to provide high utility in a task specific manner while privacy is preserved, e.g., protecting the variant coordinates genotype imputation.

ProxyTyper differs from differential privacy techniques in its approach to data protection. While differential privacy methods add noise to data to maintain privacy, this can sometimes render the data unusable due to corruption. ProxyTyper, on the other hand, uses domain knowledge to safeguard data, maintaining its usefulness while ensuring privacy. This is achieved through the creation of “proxy panels” that retain the statistical properties of the original data but anonymize subjects, alleles, and variant coordinates. This approach not only preserves privacy but also allows the proxy panels to be used as input to existing tools, ensuring high utility. This makes ProxyTyper a more practical and efficient solution for data protection, particularly in the field of genomics.

In practice, proxy panels can be used to protect genotype datasets in numerous outsourcing and collaborative tasks that require genotype level data, such as ancestry mapping, genotype imputation, and kinship estimation. To demonstrate the utility of these protection mechanisms, we focus on genotype imputation as an example task and devise an imputation outsourcing pipeline that protects the reference panel and the client’s genotype panel. In genotype imputation task, a client has a sparsely genotyped panel using, for example, genotyping arrays. The client wants to make use of the typed variants and impute the genotypes for the remaining set of “untyped” variants, which exist in a reference panel stored at another site, e.g., TOPMed panel. Most popular imputation algorithms use hidden Markov models (Li-Stephens model) and impute the untyped variant genotypes. This procedure is computationally intensive and requires sharing of the large reference panels, which may create privacy concerns. Imputation servers offer convenient services to perform imputation where client sends the typed variant genotypes to the server, which imputes the variants exclusive to the reference panel and send the results back to the client[79]. The current imputation servers share the alleles and variant coordinates in cleartext format with the client without a tunable mechanism to protect these. Furthermore, although secure imputation methods protect alleles by encryption, the variant coordinates in the reference panel are shared in cleartext. This is important because there are unexplored risks around sharing these data in reference panels: For example, the reference panels are generated from sequencing and they contain very large number of rare variants among the untyped variants and the knowledge of the coordinates for these rare variants can be directly used in a beacon-type attack to re-identify individuals, even without the knowledge of the alleles. This risk is currently not considered in many data analysis methods and data reporting policies and it may require substantial changes to how variant coordinates are reported and shared. We demonstrate that proxy panels can be used for outsourcing genotype imputation using BEAGLE without any modifications to the original imputation algorithm with only a minor decrease in imputation accuracy. Overall, our results show that proxy panel generation mechanisms can be used for protecting panels in a task specific manner against honest-but-curious entities and accidental leakages[80], which are most prevalent in terms of the adversarial model in bioinformatics. The mechanisms are flexible and can be integrated into existing pipelines for outsourcing imputation methods, can be used in AI-driven imputation methods[81,82], and in meta-imputation [83] workflows. Moreover, while we focus on imputation as our specific focus, presented mechanisms and new mechanisms can be used for other tasks such as ancestry mapping and kinship estimation.

## Results

We first present the mechanisms used by ProxyTyper framework to build proxy-panels. Next, we study the characteristics of proxy-panels, re-identification attacks, and finally present the modified imputation outsourcing protocol and its accuracy. We focus on the genotype imputation as the focus task and describe the different ways of protecting the typed and untyped variants and alleles.

### Proxy Haplotype Panel Building

We describe the protection mechanisms used by ProxyTyper for building mosaic panels at the reference and client sites (Fig. 1a), protecting the variant coordinates and genetic maps, and finally the shared alleles.

**Figure 1.**
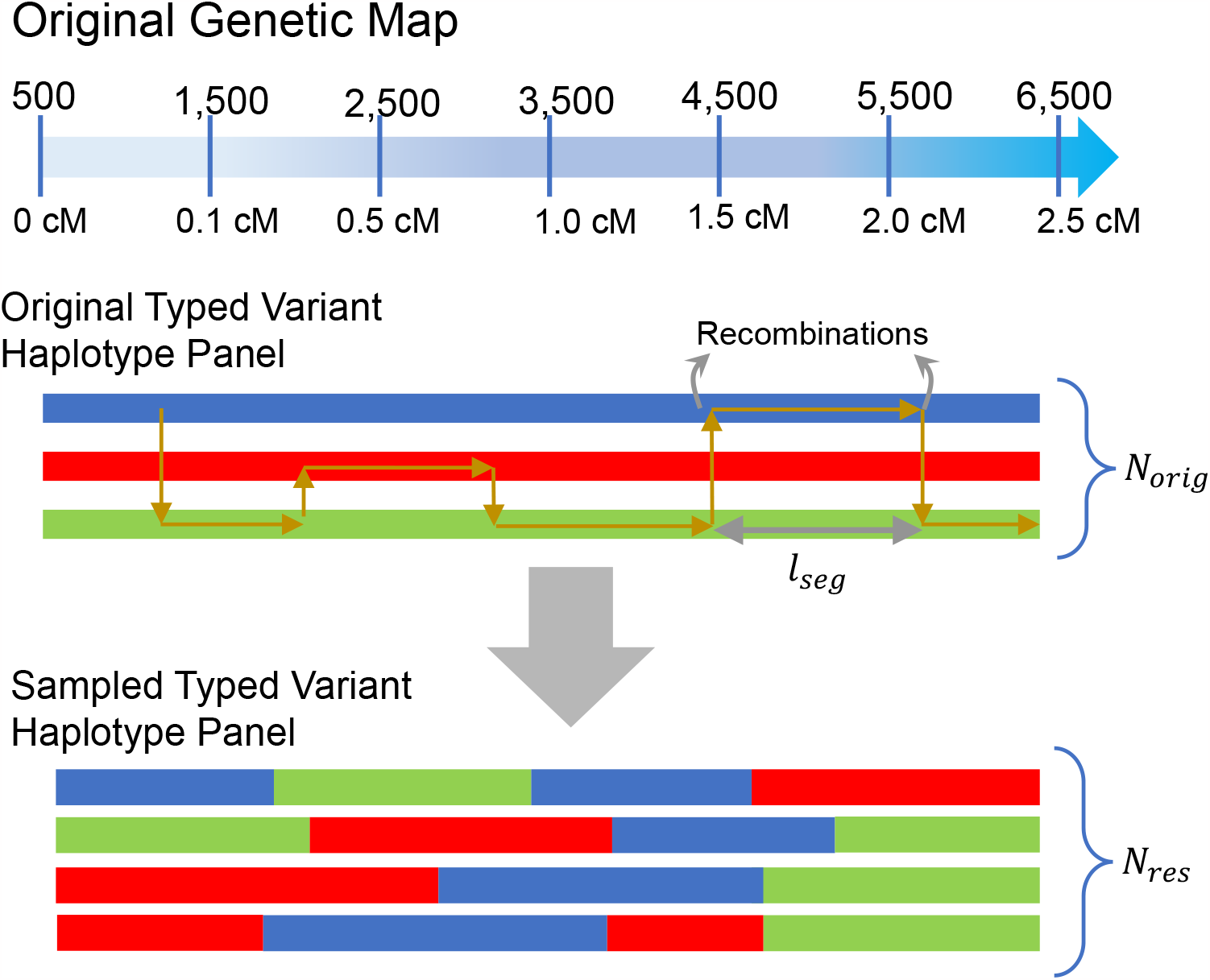
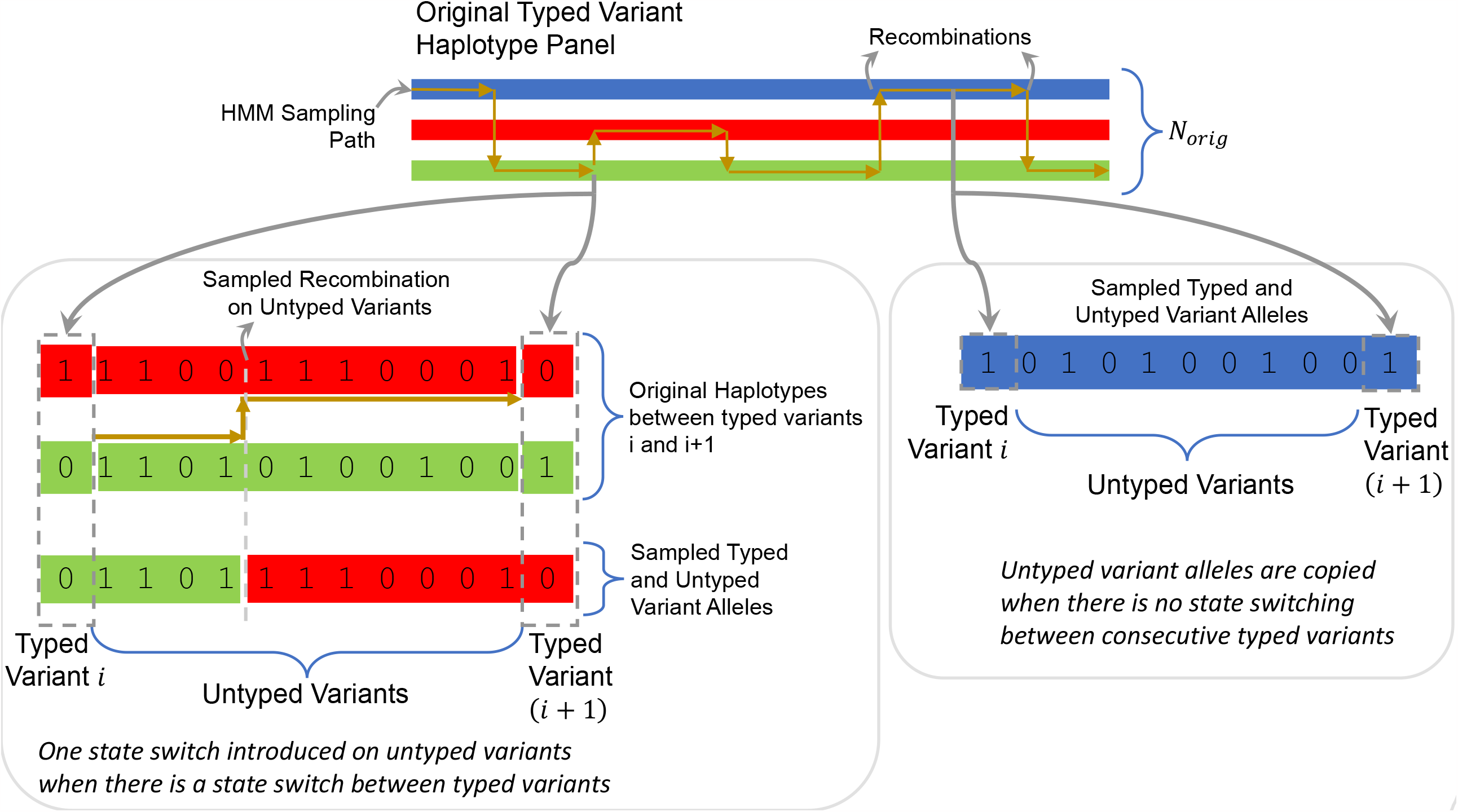
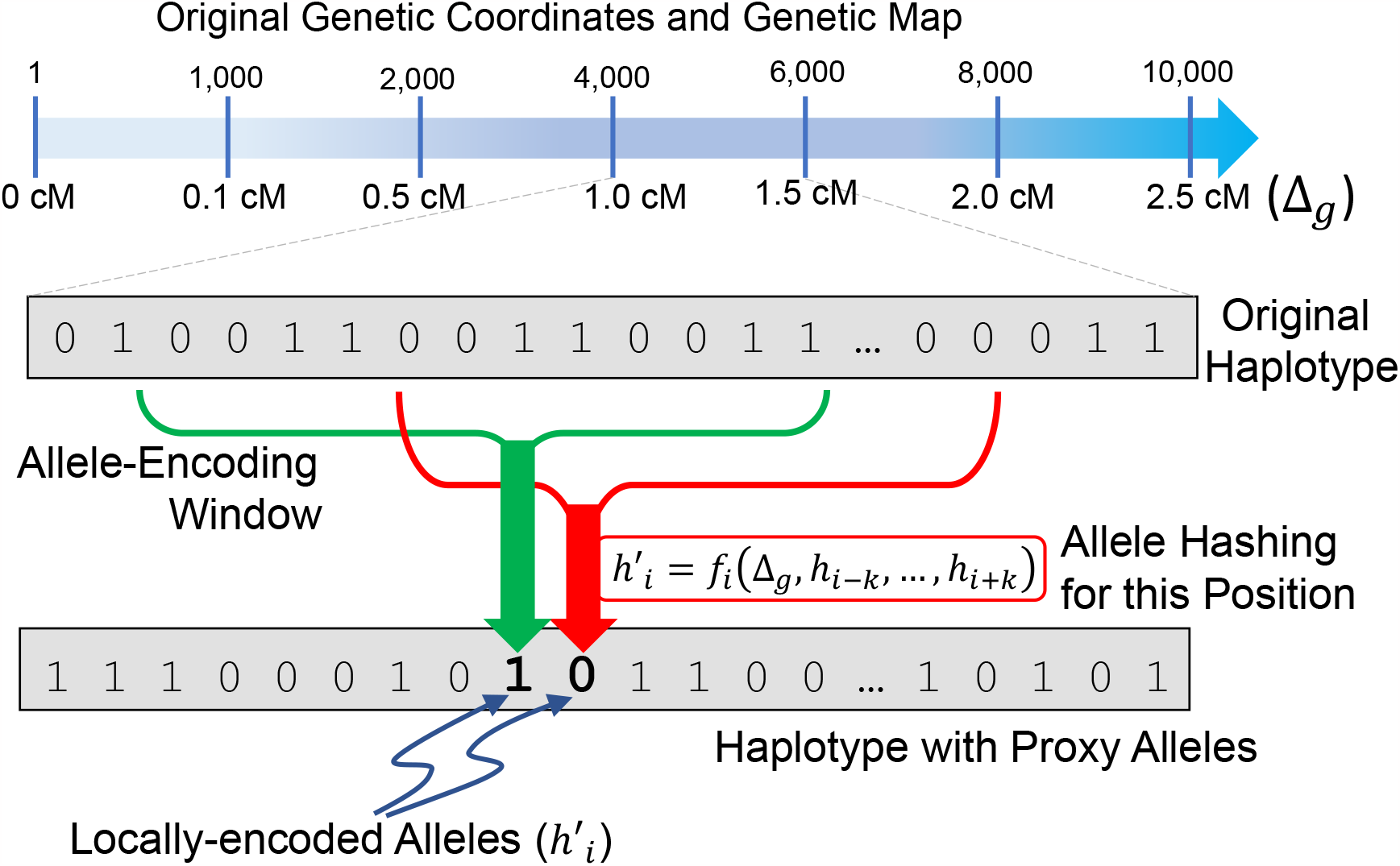
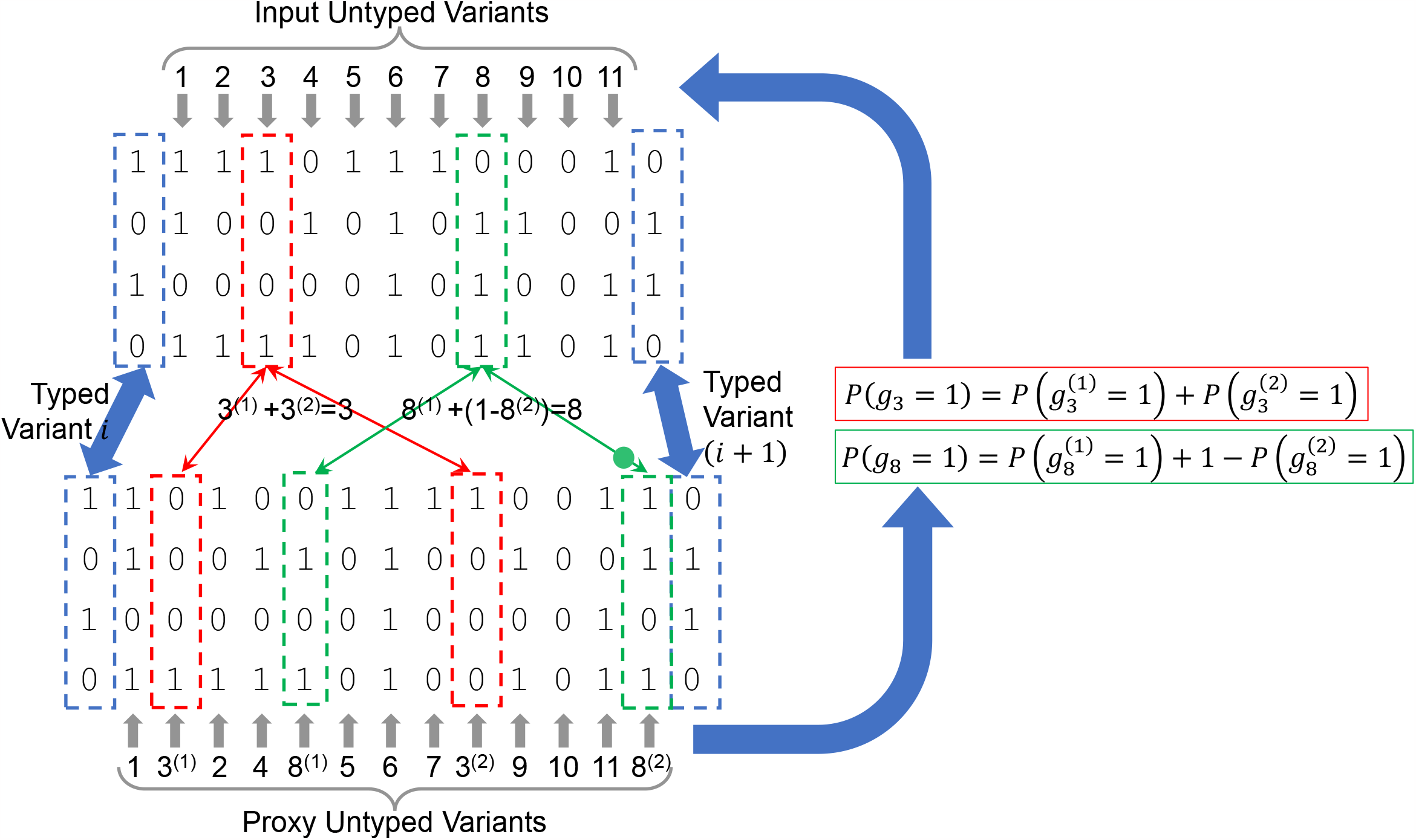
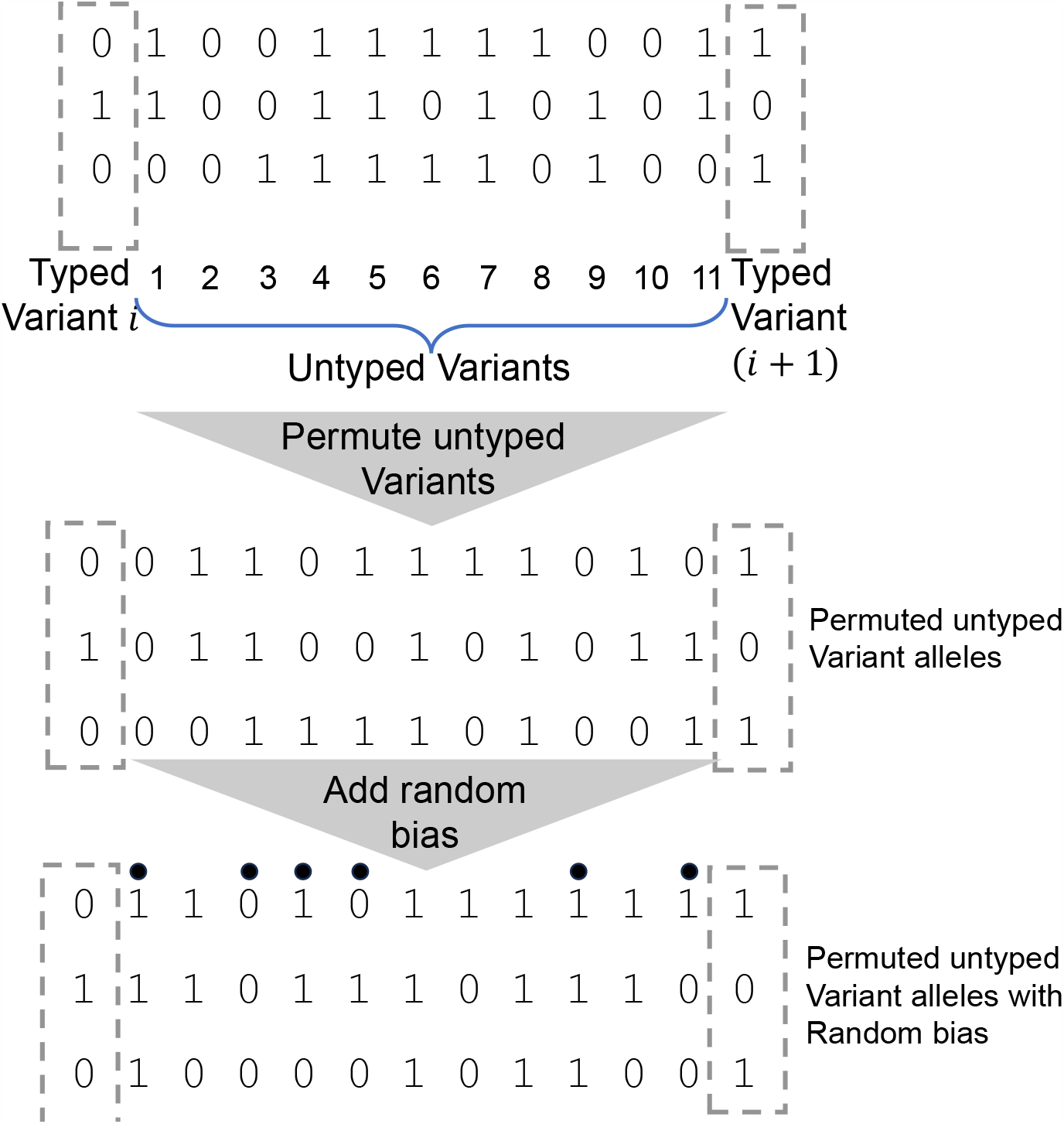
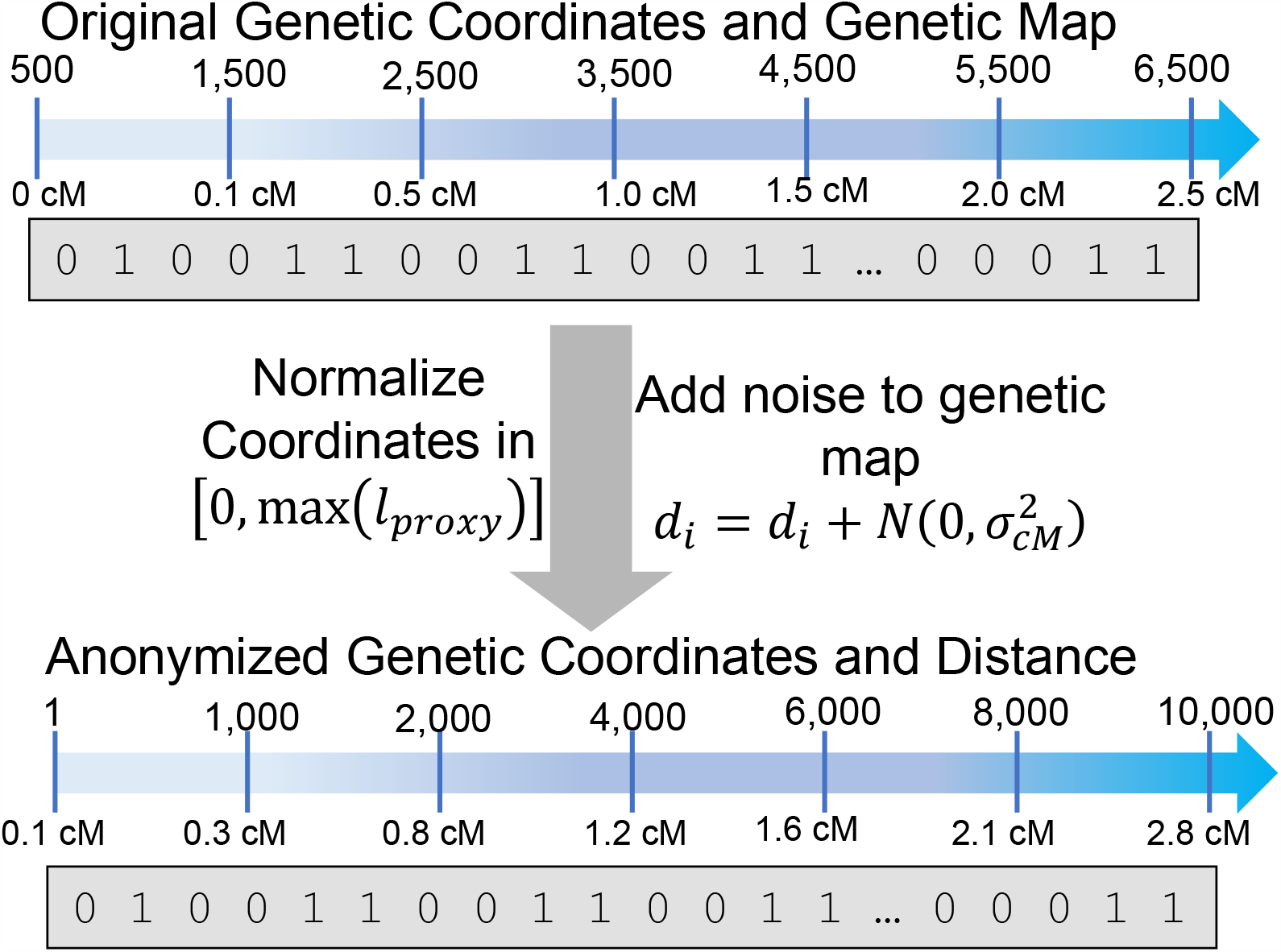
Illustration of proxy panel generation mechanisms. **(a)** Illustration of typed variant resampling. The original haplotypes are traced from left to right and recombinations (indicated by switches between haplotypes) are randomly generated using the genetic map (depicted on top). **(b)** Resampling of untyped variants following the resampling of typed variants. Given consecutive typed variants, if the sampled haplotypes are not the same (left box), ProxyTyper introduces exactly one recombination at one of the untyped variants that switches between the haplotypes assigned to the typed variants in resampling. In this example, the transition from green haplotype to red haplotype is illustrated at 4^th^ untyped variants between typed variants. If the sampled haplotypes are the same for the consecutive typed variants (right box), the alleles for the untyped variant on this haplotype is copied as the resampled untyped variant alleles. **(c)** Illustration of allele encoding (hashing). Given two windows (depicted with red and green windows), the the allele on the proxy haplotype is calculated as a combination of the alleles on the original haplotype. The two windows have independent encoding functions. The function takes the alleles and the genetic distance as parameters to calculate the hashed (encoded) alleles on the proxy haplotype (shown at the bottom). The proxy haplotype is calculated by encoding all windows using the hashes. **(d)** Protection of the variant coordinates and the genetic distance. The coordinates are normalized to a preselected value *l*_*proxy*_, which obfuscates the typed and untyped variant positions. Genetic map is obfuscated using addition of Gaussian noise with predefined variants 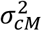. **(e)** Protection of untyped variants using (rolling) permutation between typed variant blocks. Untyped variant positions are randomly shuffled between two consecutive typed variants. **(f)** Anonymization of variant coordinates and genetic maps. The top coordinate system shows the original coordinates and genetic distance on the sensitive panel. Bottom system shows uniformly distributed coordinates and noise added genetic distances. Note that these steps do not affect the haplotype alleles.

### Mosaic Haplotypes Generation by Resampling Original Sensitive Panels

Given a panel of haplotypes, a mosaic haplotype panel is generated by a hidden Markov model (HMM) sampling of the panel. In this model, each haplotype represents a state, and transitions are performed probabilistically using modifications of Li-Stephens model[84]. Sampling starts from the first variant on the chromosome. At each variant, a new haplotype is selected and the allele on the haplotype is emitted as the sampled haplotype’s allele (Fig. 1a). After sampling all variants on the chromosome, the new haplotype is stored. Mosaic panel generation uses 2 tunable parameters for privacy-utility tradeoff: 1) Effective population size (*N*_*e*_), which tunes the average number of recombinations in sampling. 2) The maximum segment length 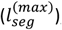, which limits the maximum length of the longest sampled continuous segment.

For the task of imputation outsourcing, ProxyTyper performs sampling only on the typed variants. The untyped variant alleles are sampled using the typed variant haplotypes (Fig. 1b, Methods). This way, the computational requirements are substantially reduced. Mosaic panel generation removes the one-to-one correspondence of subjects in query and reference panels and reduces the risks around linking subjects to external sources.

### Protection of Alleles using Hashing and Permutations

ProxyTyper uses hashing and permutation-based mechanisms to protect the alleles of typed and untyped variants for the imputation task.

#### Hashing of Typed Variants (Fig. 1c)

For protecting the identity of alleles, ProxyTyper uses a locality-sensitive hashing of the alleles. ProxyTyper replaces the original (sensitive) allele of each variant using a combination of the alleles of variants in its vicinity, i.e., locality-based hashing. This “proxy” allele protects the original allele since it is not easy to invert the proxy allele back to original allele without the knowledge of the hashing function. Given variant *i* on haplotype *j*, ProxyTyper replaces the variant’s allele with an allele calculated by a modulo-2 hash function of the alleles for variants in the vicinity of variant *i*. To introduce randomness, ProxyTyper uses a function that combines upto 3^rd^ degree combinations of the alleles of variants in the vicinity:

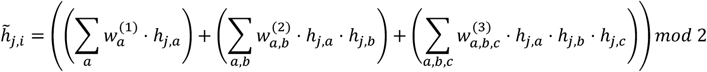

where 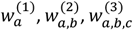 denote the randomly selected variant-specific weights for calculating the hash for 1^st^, 2^nd^, and 3^rd^ degree combinations for the variants at *a, b, c*. To decrease computational complexity, the variants are constrained to a maximum distance, i.e., |*i* − *a*| < *n*_*vic*_. The basic idea of hashing is to spread the allelic information within a window since each variant’s allele randomly contributes proxy alleles within the hashing windows. For tasks like imputation, the obfuscated alleles are expected to preserve the haplotype information while protecting the identity of the original allele.

#### Distance-dependent Random Selection of Hashing Weights

At each position, the weights for hash are selected randomly to ensure that the proxy alleles are generated independently without leaking model information (Methods). Given a small window, the weights are selected randomly using genetic distance dependent manner to ensure that the probability of a hashing window does not cross recombination hotspots. More formally, given two variants *i*_1_ and *i*_2_, a weight is assigned to 1 with probability 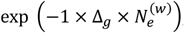, where 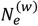 is a tunable parameter for selecting weights where Δ *g* denotes the genetic distance (in centiMorgans) between the variants.

The final “proxy haplotypes” are calculated by evaluating the hashing function at each variant of each haplotype. Due to modulo arithmetic, the proxy alleles are binary-valued, and they look like valid alleles albeit with virtually no correlation to the original (sensitive) alleles. The proxy haplotypes provide utility for tasks such as imputation because they preserve local haplotype frequencies between two panels that are hashed using the same hashing functions. This relationship is directly related to the vicinity size *n*_*vic*_, i.e., smaller *n*_*vic*_ provides better match to original haplotype frequencies. On the other hand, larger *n*_*vic*_ provides better protection at the cost of decreasing the concordance to original haplotype frequencies.

#### Partitioning and Permutation of Untyped Variants (Fig. 1d, 1e)

Hashing of untyped variants may add extensive computational burden because untyped variants are much larger in size than the typed variants. ProxyTyper instead uses a partitioning and permutation mechanism to protect the untyped variants. Given an untyped variant *i* that resides between typed variants *a* and *b*, the haplotypes that harbor the alternate allele of the untyped variant are randomly separated into two sets. Next, ProxyTyper generates two proxy untyped variants at random positions *i*_1_ and *i*_2_ between *a* and *b* (Fig. 1d). The alleles in the first haplotype set are assigned to *i*_1_ and those in second set of haplotypes are assigned to *i*_2_. This procedure effectively partitions the alleles of the untyped variants into two new proxy untyped variants such that the allele frequency of original untyped variant is the sum of the two proxy-variant allele frequencies.

ProxyTyper also performs a final shuffling and inversion of the proxy untyped variants within a sliding window, i.e., a permutation filter (Fig. 1e). The permutation is constrained to be among typed variants to conserve the LD information between the typed and untyped variants for accuracy of imputation. This is justified since the imputation tools execute HMMs on the typed variants and imputation of untyped variants rely on the typed variant. The reference allele for each proxy untyped variant is further randomly flipped with 50% chance (Fig. 1d, 1e). The final proxy allele frequencies do not exactly reflect the allele frequencies of the original panel but can be still useful for tasks such as genotype imputation where local haplotype information is sufficient for performing the task.

### Protection of Variant positions and genetic maps

Homer[38] and LRT-type re-identification attacks[39] attacks require matching the variant positions between the reference panel and the mixture panel so that attack statistics can be calculated for identifying a target individual with known genotypes. To ensure that the proxy variant positions are not directly linkable to external panels, ProxyTyper obfuscates the variant positions by replacing variant positions to be distributed uniformly on a predefined range (Fig. 1f). Even though the variant positions are obfuscated, genetic maps can be aligned to public sources to match them and reveal exact locations. To protect the genetic maps that are shared among the sites, ProxyTyper perturbs the maps by noise addition with a user-specified deviation *σ*_*map*_, measured in centiMorgans (cM). For example, given *σ*_*map*_ = 0.05 *cM*, corresponds to approximately 50 kilobases (1 megabase/cM) of perturbation to the position of the variant. Of note, for the imputation panels, ProxyTyper generates the anonymized genetic maps only for the typed variants because untyped variant genetic maps are interpolated by BEAGLE from typed variants. It should be noted that imputation software such as BEAGLE does not require variant coordinates when genetic maps are available. This is expected since imputation HMMs work on executing Li-Stephens model using the genetic distances and coordinates (in base pairs) are not needed for accurate imputation.

### Comparison of Proxy Panel with Original Panel

In this section, we compare the proxy panel and the original panel with respect to their simple statistics. Since entities would have access to only the proxy panel, the comparisons presented here would not be possible by an adversarial honest-but-curious entity, which is the main adversarial model we are focusing on. We, however, compare the original panel and proxy panels to highlight how well the proxy panels reflect the original panels that they were derived from.

Each haplotype in the proxy haplotype panel is generated by 1) resampling, 2) hashing of the typed variants alleles and 3) partitioning and permutation of the untyped variants (only for the reference site), 4) Protection of the variant positions and genetic maps. We first asked whether the proxy haplotypes leak information about the original panel at the level of variants. We compared the proxy panel and the original panel in terms of genotype and variant-variant linkages.

#### Comparison of genotypes and haplotype frequencies in original and proxy panels

We first assessed the correlation between the original variant genotypes. We extracted the genotypes of 661 individuals in AFR population of the 1000 Genomes Project and divided the panel into 2 subpanels comprising 330 and 331 subjects. We used the first panel (original panel) and generated the corresponding proxy panel with the same number of subjects. The second panel is used as a matching control (holdout) panel. For each variant, we correlated the original panel genotypes (alternate allele dosage) with the genotypes of the proxy panel among individuals. We observed no significant correlation between the genotypes of the original and the proxy panel (Fig. 2a). Further, the genotype correlations of original-vs-proxy did not exhibit any enrichment compared to the correlations of original-vs-holdout panel. This is expected because the resampling step breaks one-to-one correspondence between subjects. When we remove resampling step in proxy panel generation (Fig. 2a), the distribution of correlations exhibits a large variance. This is also expected since for some variants, the hashing may involve a small number of surrounding variants due to random hashing weight selection (Methods) which leads to high correlation to the original variant alleles.

**Figure 2.**
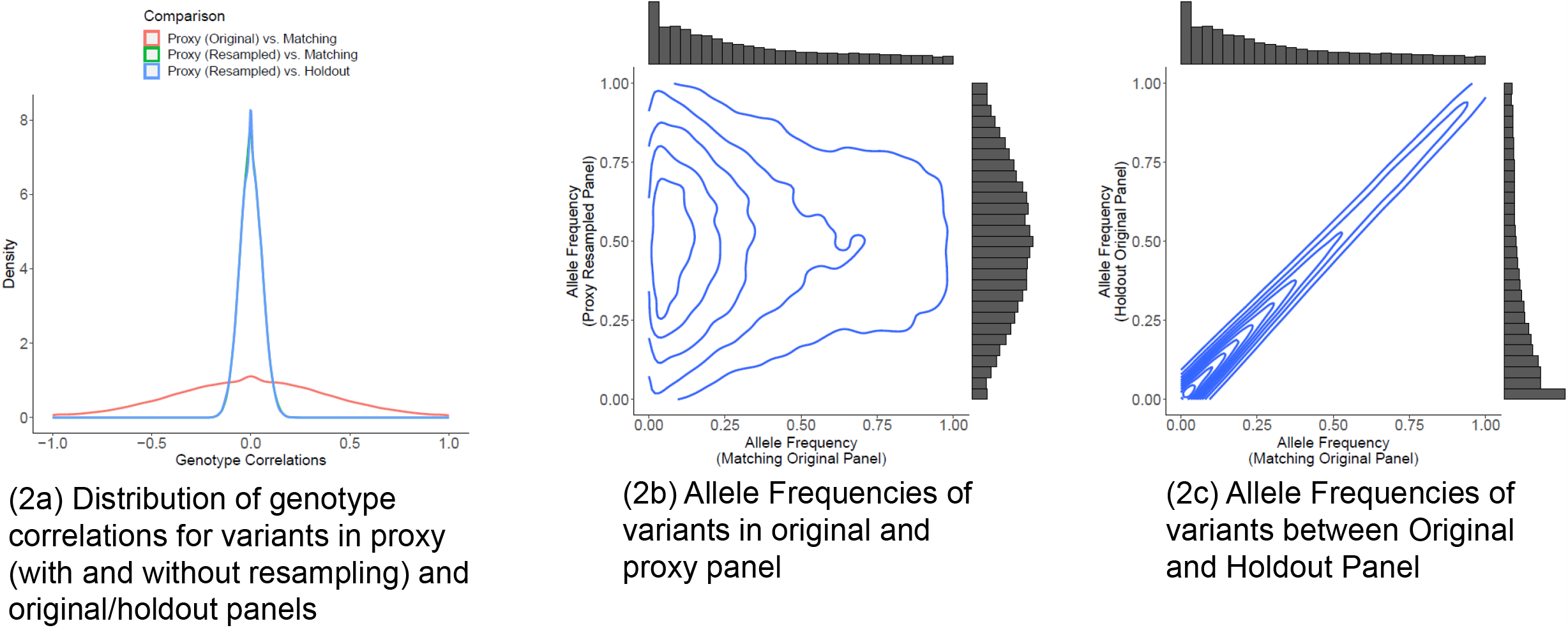
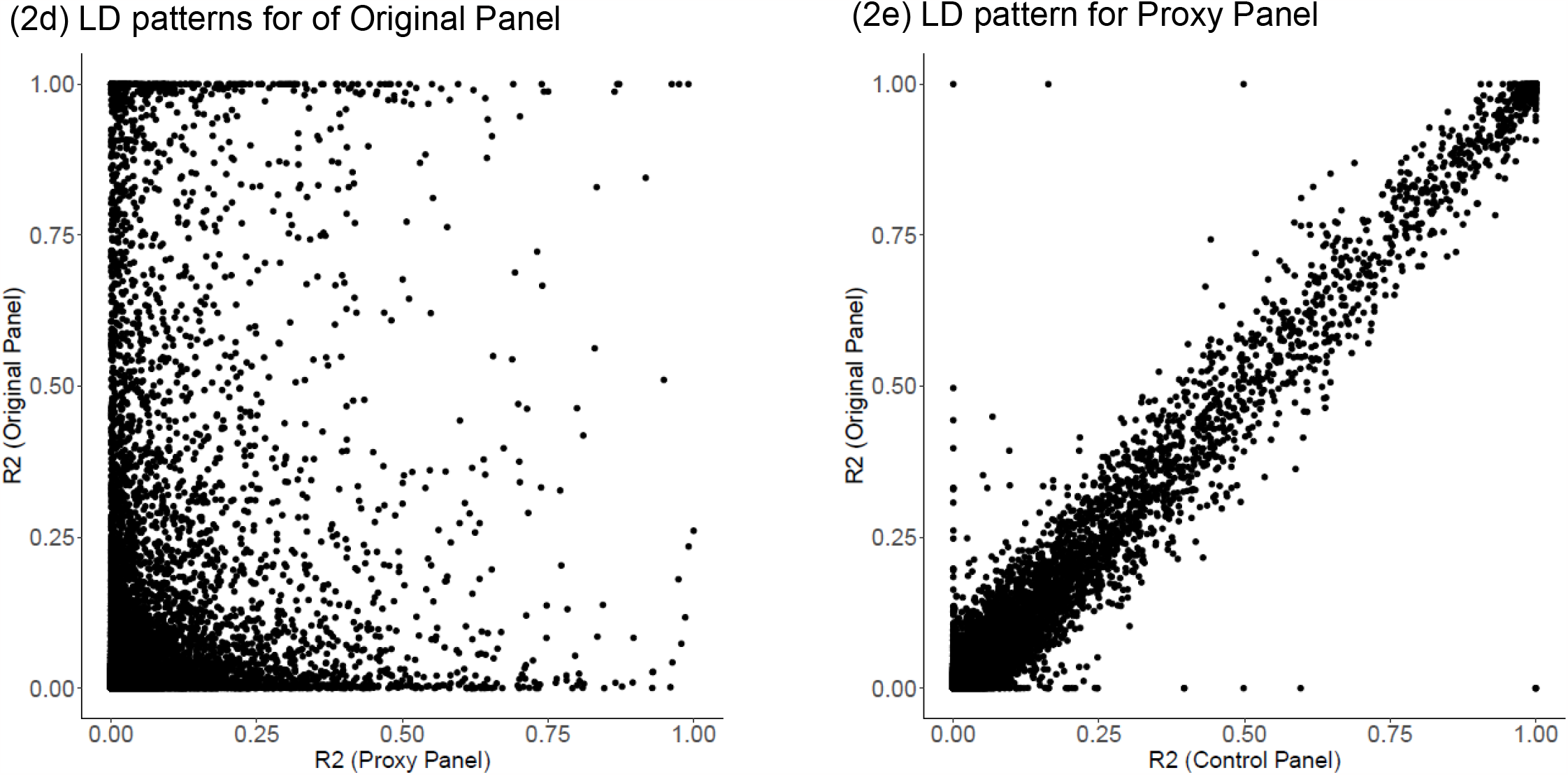
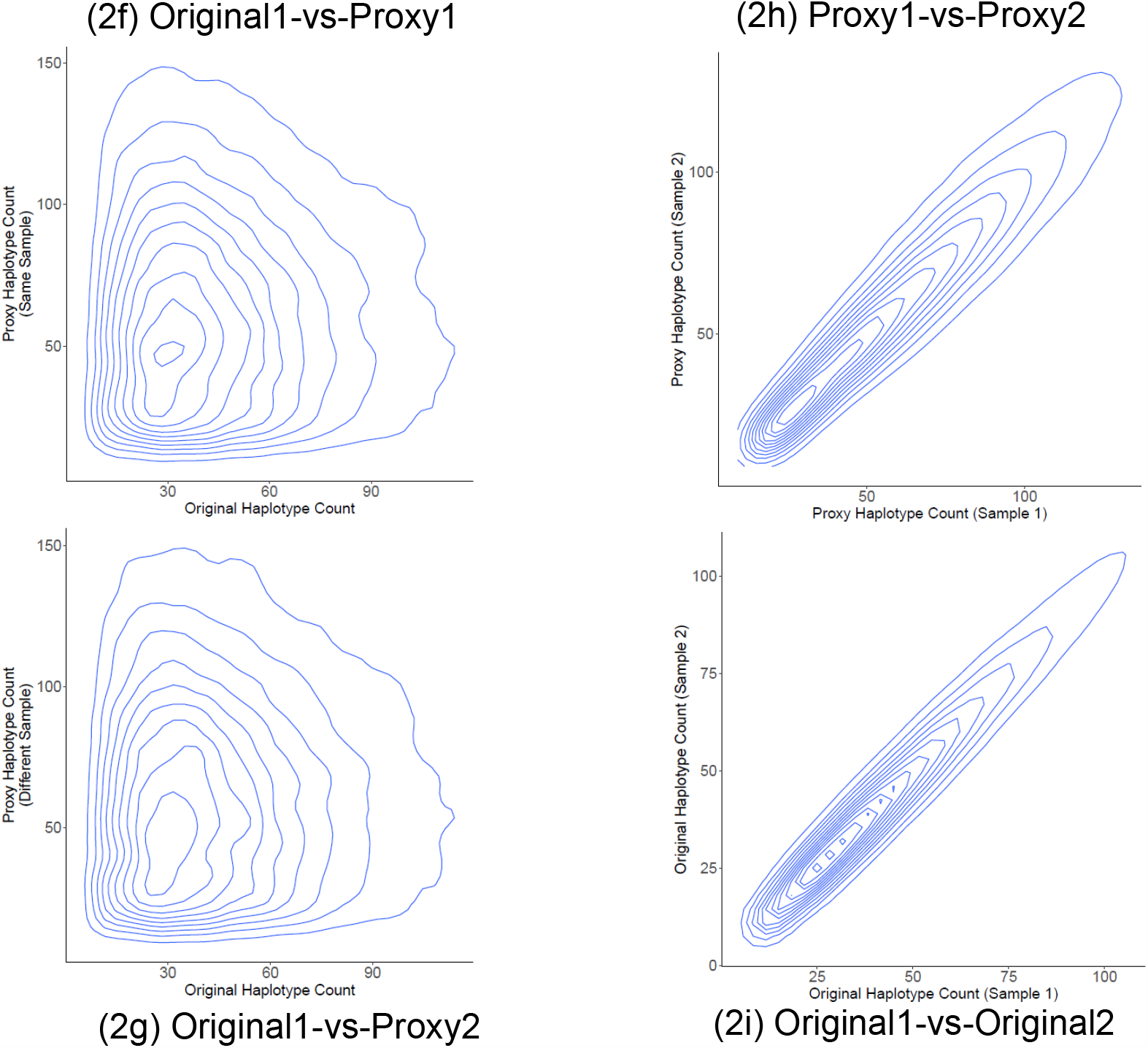
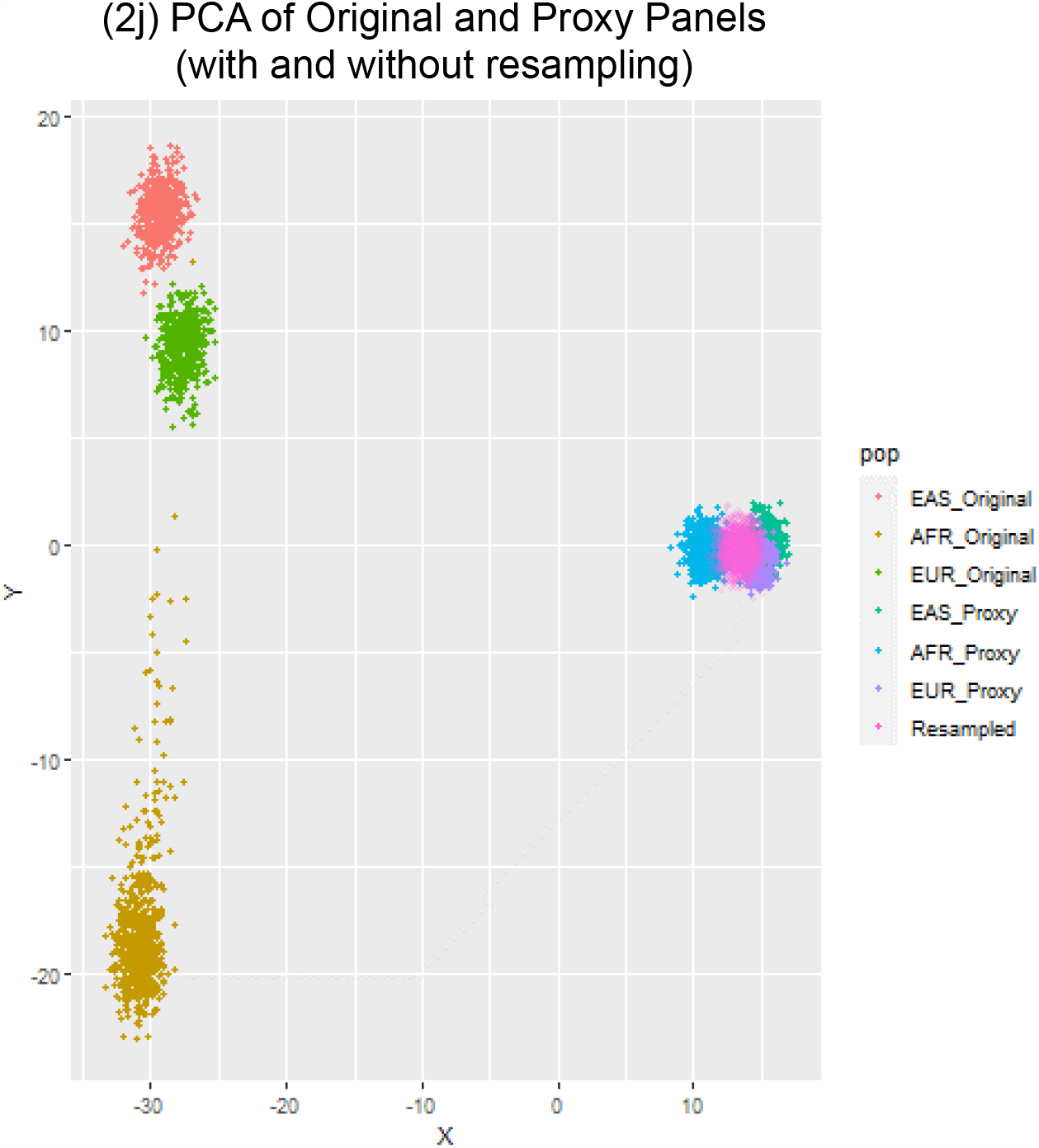
Comparison of proxy and original panels. **(a)** Distribution of Pearson correlation of genotypes between proxy (with and without resampling), matched and holdout panels. **(b)** Density plot shows the distribution of allele frequencies in proxy panel (y-axis) versus the original panel (x-axis). **(c)** Density plot shows the distribution of allele frequencies in holdout panel (y-axis) versus the original panel (x-axis). **(d)** Distribution of the variant-variant correlations in 500 variant windows that illustrates linkage disequilibrium patterns in the proxy panel and the original panel for matching pairs of variants. Each dot corresponds to a pair of variants whose R2 correlation is shown on x and y axes on proxy and original panels, respectively. **(e)** The comparison of variant-variant R2 in cleartext original and holdout (control) panels. **(f)** The distribution of local k-mer counts in proxy panel and matching original (original1) panel. **(g)** The distribution of local k-mer counts in proxy panel from proxy panel generated from original2 panel and the cleartext original1 panel. **(h)** The distribution of local k-mer counts in proxy panels generated from original1 and the original2 panels. **(i)** The distribution of local k-mer counts in cleartext original1 panel and the cleartext original2 panel. **(j)** Distribution of the subjects in original panel and the proxy panels from Principal component analysis (PCA). The first two components are shown in the scatter plot. Each dot is a sample and colors indicate the populations for the corresponding individuals. Proxy panels are generated with and without resampling indicated by colors. Note that when resampling is used, population is not depicted since each haplotype is a mosaic of all populations.

We next compared the allele frequency of each variant in the original and proxy panels. We observed there is virtually no correlation between allele frequencies of variants in the original and proxy panels (Fig. 2b). As expected, holdout panel exhibits very high concordance to the original panel in allele frequencies. (Fig. 2c)

#### Variant-variant Linkage and Local Haplotype Frequency Concordance between Original and Proxy Panels

We next calculated the Pearson R2 coefficient between genotypes of variant pairs in a sliding window of 500 variants to estimate the linkage patterns in original and proxy panels. We observed that the variant-variant correlation patterns do not exhibit clear concordance although there is significant correlation of (Pearson R=0.33) (Fig. 2d). It should be noted that the linkage concordance between the original and control panels is much clearer (Fig. 2e) (Pearson R=0.99). We next compared the local haplotype counts between the original, proxy, and the control panels by extracting the k-mers (k=11) in sliding windows and calculating the number of unique k-mers in each window. Although there is a small correlation between the number of unique k-mers at each position of the proxy and original panels (Pearson R=0.01), the concordance is much weaker (Fig. 2f, 2g) than the two matching panels (Fig. 2h, 2i). In particular, we observed that the proxy panel harbor an excess of unique k-mers per position compared to the original panel. This result demonstrates that random hashing increases the entropy of the k-mer (local haplotype) distribution at each position.

#### Population Stratification of Proxy-Panels

We finally performed Principal Component Analysis (PCA) using the proxy panel genotypes to evaluate how well the population-level information is reflected in proxy panels in comparison to the original panel. In this analysis, we first extracted the genotypes of the 1000 Genomes Project from the EUR, AFR, and EAS populations and performed PCA on the original genotypes and the proxy genotypes. When the sampling step is performed, PCA did not provide any information about the ancestry of the proxy panels (Fig. 2j). When we generated the proxy panel without the resampling step, the proxy panel separated into 3 distinct clusters. This is expected because sampling shuffles the haplotypes into mosaics and ancestry information is lost among the sampled mosaic haplotypes. To assess if it is possible to match the subjects to each other when sampling is not used, we aligned the SNPs by their chromosomal order and we merged proxy and original genotype matrices into a pooled genotype matrix and performed PCA on the pooled matrix. This shows that the original and proxy panels separate into 6 distinct clusters (Each corresponding to an ancestral group in original and proxy panels) with no clear mixing between them.

Overall, these results indicate that proxy panels do not leak substantial first order (allele frequencies) or second order information (variant linkage or population stratification) about the original sensitive panels and it is unlikely to perform re-identification using proxy panels without a malicious intent (e.g., stealing hashing parameters). Application of Homer’s t-statistic and LRT attacks are not immediately possible on proxy panels since variant coordinates are obfuscated and the genotypes are hashed. We also hypothesize that linking attacks (matching each subject in proxy panel to an external panel) are unlikely when sampling step is used because each proxy-haplotype is a random mosaic of the original haplotypes.

### Viterbi Decoding of Proxy Alleles and Re-identification Attacks

Our previous results indicate that there is small but significant correlation linkage of consecutive variants in original and proxy panels (Fig. 2). The main risk for re-identification in the proxy-panels will stem from decoding of the proxy-alleles to infer the original panel by matching the haplotype frequencies using an external panel, followed by a re-identification attack such as an LRT attack. This requires an adversary to decode the hash function weights at every variant, which is unlikely without external knowledge (e.g., stolen or leaked parameters which may lead to *known-plaintext attacks*). We assume that the adversary only has access to the proxy panel haplotype matrix with some knowledge of the hashing function such as the size of vicinity window but does not know the weights of hashing. While it is not immediately available, the size of vicinity window used for hashing could potentially be inferred from the proxy panel’s k-mer frequency statistics.

One attack for decoding the proxy panels can make use of two properties of the hash function: Firstly, hashing of consecutive variants use overlapping windows, i.e., if the adversary can guess the original k-mers at a position, these k-mers can be used to make a better decoding prediction for the next variant. Secondly, the k-mer frequencies between the proxy panel and original panel are somewhat concordant, e.g., a high frequency k-mer in the original panel is likely a high frequency k-mer in proxy panel and vice versa. Thus, the adversary does not have to infer the weights of the hash function but try to match the k-mers based on their similarity in proxy and reference frequencies to decode and map the proxy k-mers back to original k-mers while using the k-mer at previous position as a constraint and perform a *frequency analysis attack*[85]. This process can be implemented in an HMM (similar to genotype imputation) while considering the recombination rates (estimated from genetic distances) as further constraints on the decoded alleles.

To test the viability of this attack, we implemented a modified version of the Li-Stephens hidden Markov model that can decode the k-mers in proxy haplotypes by comparing their frequencies in the proxy panel to those in a separate reference dataset (Methods). While decoding a proxy haplotype, HMM starts from the first variant. At each variant on the proxy haplotype, HMM constrains the possible reference k-mers in comparison with the previous position, and the frequency concordance between the proxy haplotype’s k-mers and the original k-mers, and integrates the recombination frequencies (i.e. haplotype switch) using the genetic distances. Finally, it assigns a score to each reference haplotype that quantifies the likelihood that the original alleles are emitted from the respective reference haplotype. After all haplotypes are scored for all variants, Viterbi decoding is used to trace back the highest scoring decoded allele sequence for the proxy haplotype. Given that each variant is independently hashed, this procedure will find the haplotypes with largest joint probabilistic score that combines the haplotype frequency concordance between the proxy panel and the reference panel, and the recombination rates of consecutive SNPs.

Of note, the computational resources for executing this HMM is at least as large as a Viterbi-based imputation HMM. Furthermore, the state reduction techniques[86] are not directly applicable to the proxy-decoding HMM because the state-switching probabilities rely on both the variant location and the k-mer concordance. Therefore, unlike Homer’s attack and LRT attacks, which can be easily performed by honest-but-curious entities, proxy-decoding attacks require large computational resources with a malicious intent.

To test accuracy of the re-identification using decoded proxy panels, we divided the 661 subjects in AFR population of the 1000 Genomes Project into three panels. The first panel (100 subjects) is used to generate a proxy panel. The second panel (461 subjects) is used as the reference panel for proxy-decoding and LRT attack. The third panel (100 subjects) is held out as a control for the baseline LRT attack on non-matching individuals (control panel). We focused on the 87,960 variants on chromosome 1. Since the variants used by the proxy panels are obfuscated, we divided the variants into two categories by assigning every other variant to the proxy panel and the remaining variants to the reference panel such that the panels contained 43,980 non-overlapping variants. This way, we assume that the attacker approximately guesses the positions of the proxy panel variants. To decrease computational requirements, we focused on the variants between 10-100 million base pairs, which results in 19,379 variants. We generated the proxy panel for the first panel and decoded the proxy-haplotypes using the reference panel. We next evaluated the accuracy of the decoded alleles (for proxy panel variants) by calculating the concordance of the known original panel alleles and the decoded alleles. As a baseline control, we shuffled the decoded panel subjects and calculated the average allelic difference between shuffled decoded panel and the first panel. Overall, the proxy-decoding procedure does provide improvement of around 7% over control (30% vs 23%) (Fig. 3a) when small hashing window length is used (hashing with 7-mers). As the hashing window length is increased to 17-mers, the accuracy of decoded proxy is 3% lower than control (30% vs 27%) (Fig. 3a).

**Figure 3.**
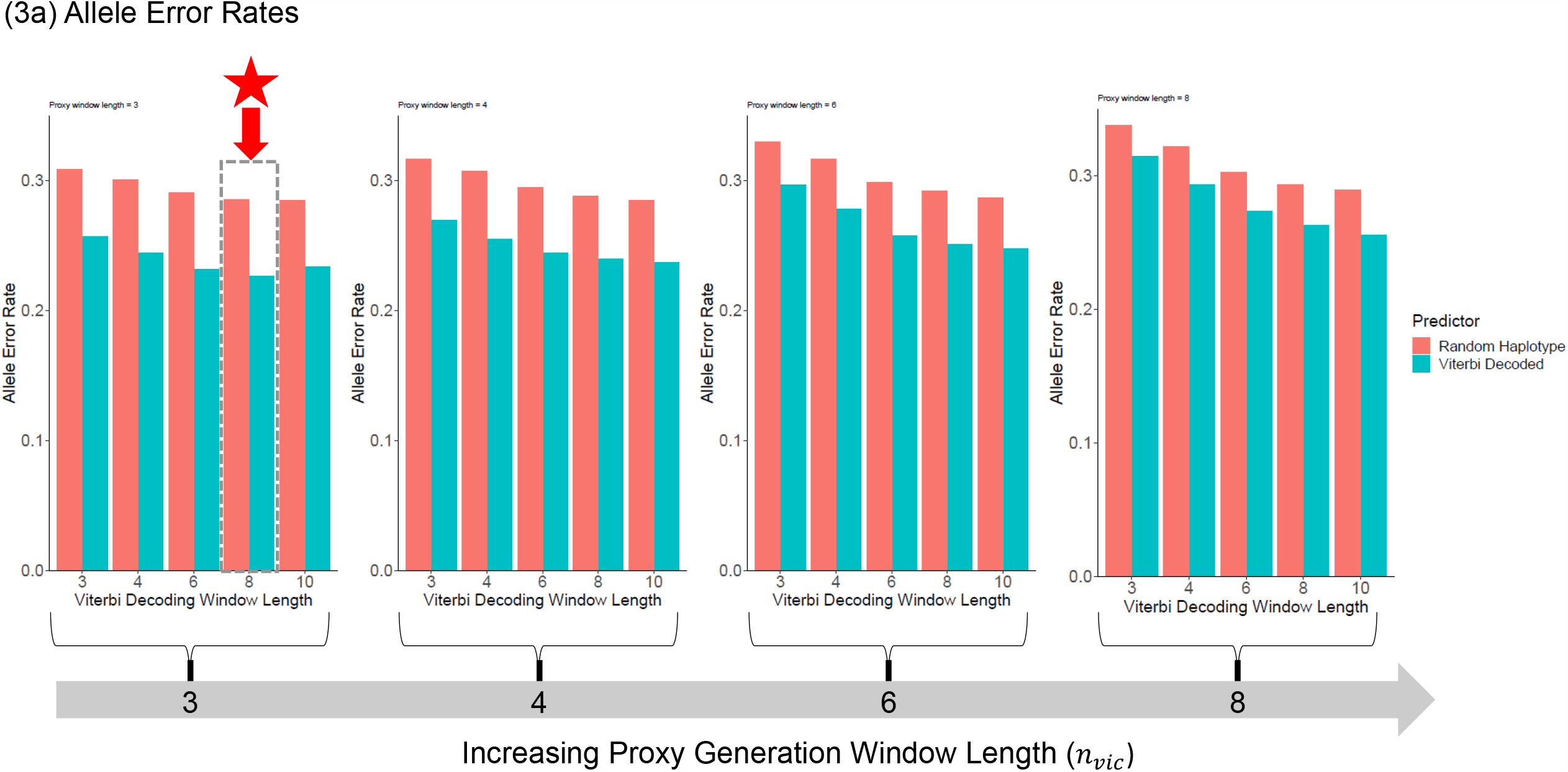
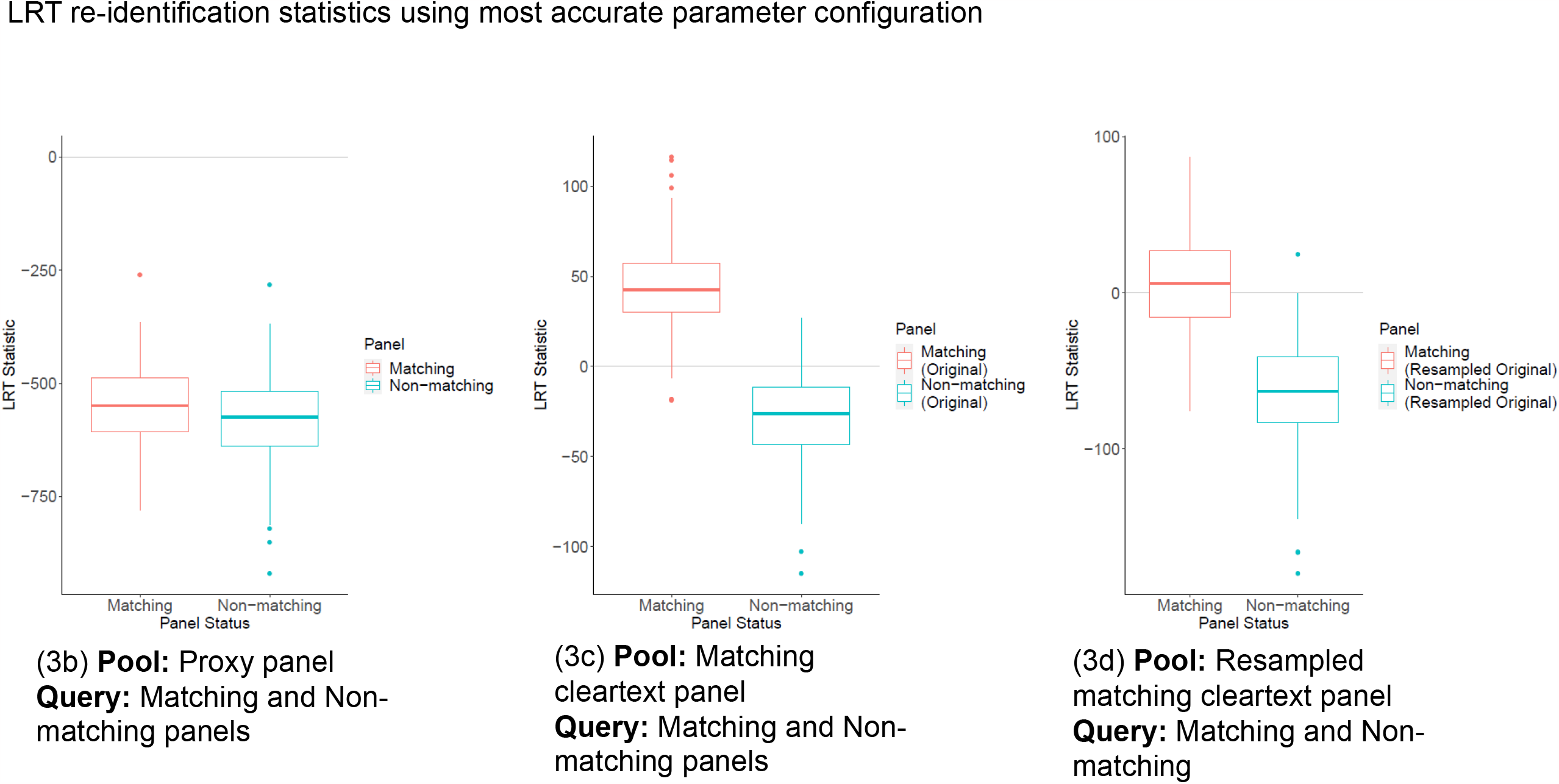
Viterbi Decoding and LRT attack on proxy panels. **(a)** Allele decoding accuracy. Each plot shows the allele error fraction with increasing window length used for the decoding. In each plot, bar colors indicate the error for Viterbi (Blue) and error rate for the randomly matched haplotypes (Red). Between different plots, the hashing allele hashing window length is increased from 7 to 17-mers indicated on the horizontal arrow at the bottom. Note that the vicinity variant size for hashing corresponding to these parameters varies from 3 to 8. **(b)** Distribution of Sankararaman’s LRT statistic using matching panel (Red) and non-matching panel (Blue) as the query individuals and the decoded panel as the pool. **(c)** Distribution of LRT statistics using matching panel (Red) and non-matching panel (Blue) as the query individuals and the original matching panel as the pool panel (No proxy panel). This is used as the baseline re-identification scenario when pool panel’s allele frequencies are exactly known. **(d)** Distribution of LRT statistics matching panel (Red) and non-matching panel (Blue) as the query individuals and the resampled matching panel as the pool without hashing of variants.

#### LRT Attack using Decoded Proxy Panel

We next used the proxy decoded alleles of the proxy panel in LRT-based re-identification attack. For this, we used the decoded proxy panel as a pool in which the adversary tries to re-identify an individual whose genome is available. The reference panel (461 subjects) used for decoding is again used as the reference panel for the LRT-based attack. For each original panel subject, we calculate the LRT statistic. As a control to individuals in the pool (original panel), we calculated the LRT statistic for the subjects in the held-out control panel (100 subjects), who did not participate in the pool. When the LRT statistics are compared, we did not find a clear separation between the distribution of LRT scores for the original panel subjects and the control panel subjects (Fig. 3b). When we applied LRT on the cleartext panels (no proxy panels), the test statistic clearly distinguishes the original and control panels for re-identifiability (Fig. 3c). We also calculated LRT test on the sampled original panel without hashing, i.e., usage of mosaic haplotypes only and observed that original panel can be easily distinguished from control panel (Fig. 3d). However, the test may be miscalibrated (LRT statistic is below 0 for approximately half of the original panel subjects). This result demonstrates the importance of using the allele hashing while sharing haplotype datasets. We also observed that when the population of the reference panel in LRT attack does not match to the ancestry of the proxy panel, it is challenging to calibrate LRT attack to re-identify individuals. This is concordant with previous studies that highlighted the importance of matching ancestry of the reference panel to the target individual and the pool that is being tested[52,53]. Unless the adversary has the knowledge of exact ancestral groups of the individuals in proxy panel, we expect that the re-identification will be more challenging than our approach. Notably, our previous results show that a PCA analysis does not immediately reveal ancestral status of subjects in the proxy panel (Fig. 2k). This renders it more challenging for an adversary to build a reference panel for re-identification purposes.

### Usage of Proxy Panels in Outsourcing of Genotype Imputation

We first describe the basic idea for using proxy panels in genotype imputation outsourcing services, e.g., imputation servers. These servers store large reference panels (e.g., TOPMed) and provide the imputation-as-a-service for researchers. The researchers upload their genetic data consisting of sparsely genotyped (e.g., genotyping arrays) study subjects to the server. The server uses the reference panel and runs the imputation software (e.g., BEAGLE, Minimac4) where overlapping typed variants are used for executing the hidden Markov model to impute the genotypes for variants in the reference panel that are untyped in client’s variants. The final results are returned to the researcher. This process requires the researcher to submit the genomes in cleartext format (i.e., unsecure), which may create privacy concerns or may not even be allowed by data usage agreements.

To make use of proxy panels in the imputation service, the client and server execute a protocol that we describe below (Fig. 4a):

**Figure 4.**
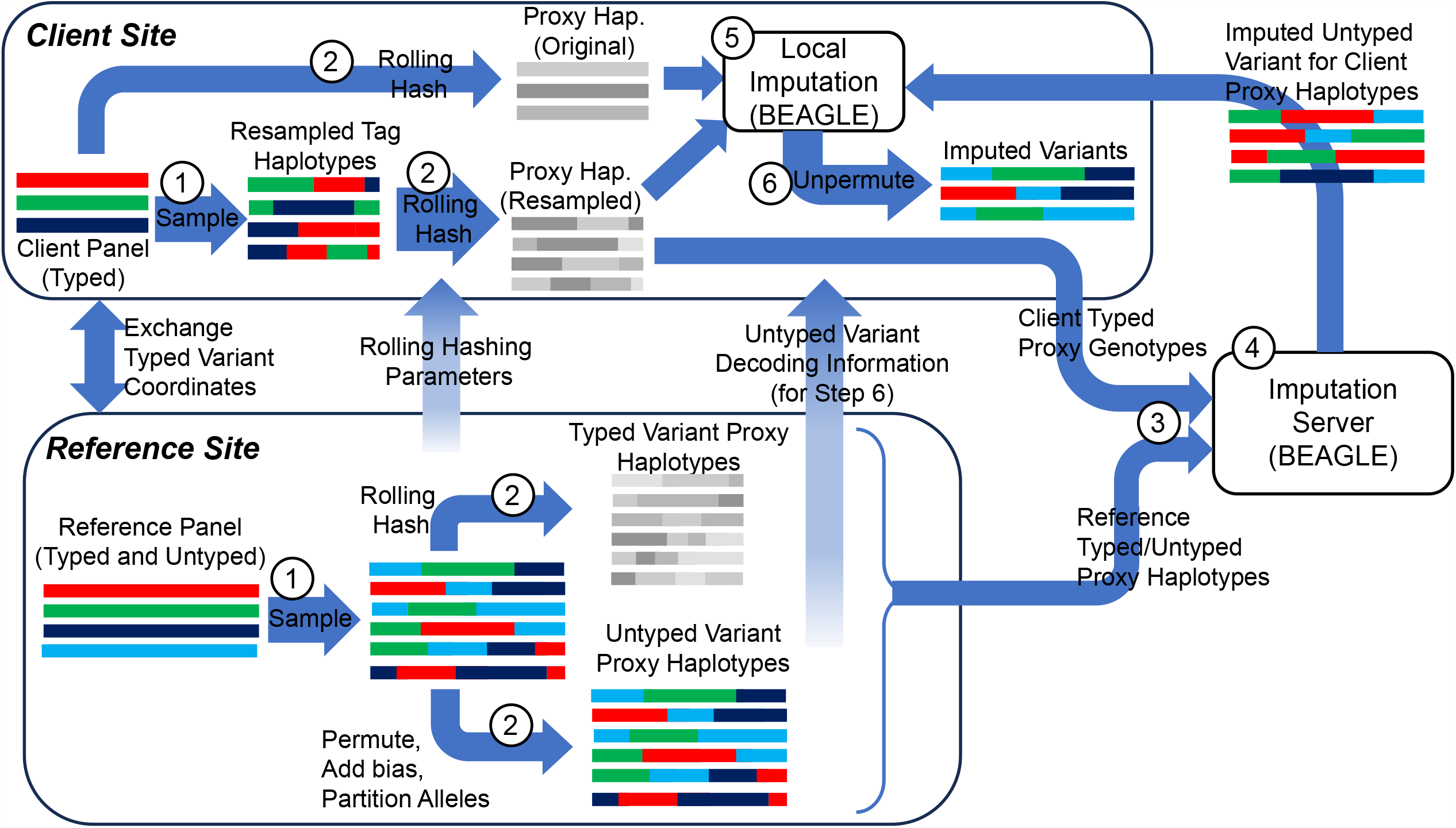
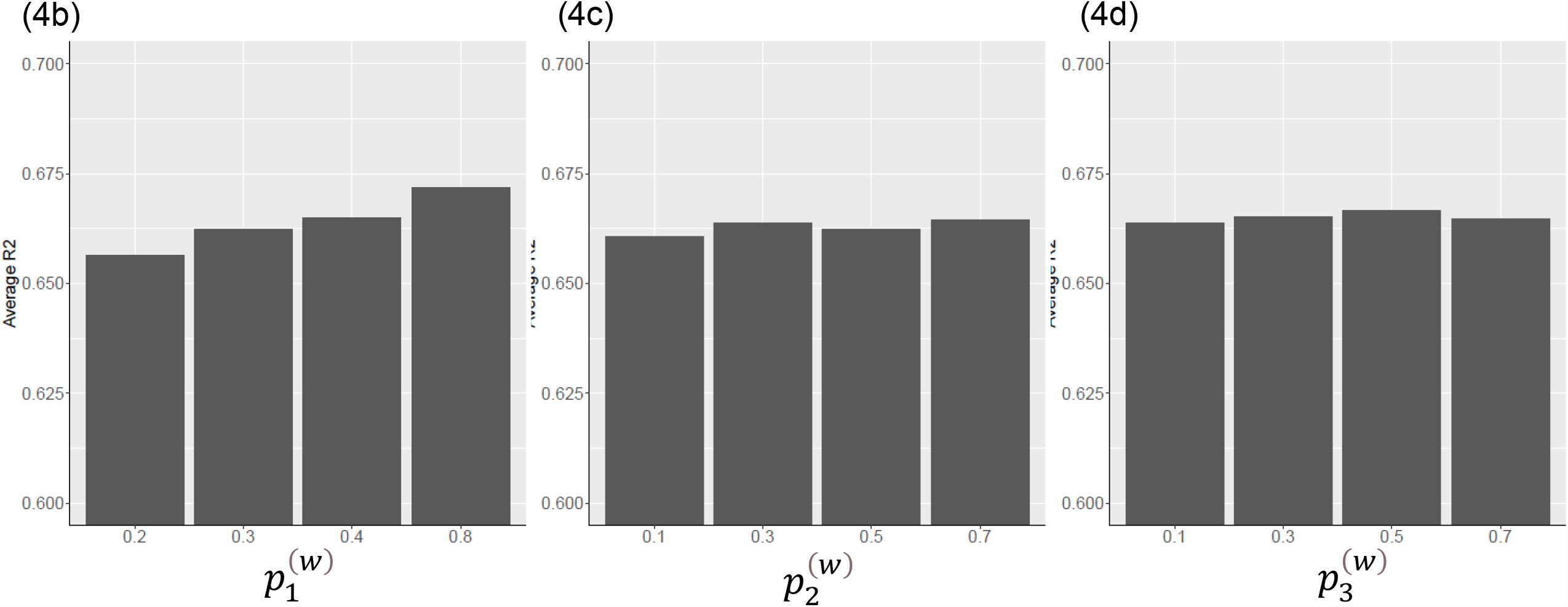
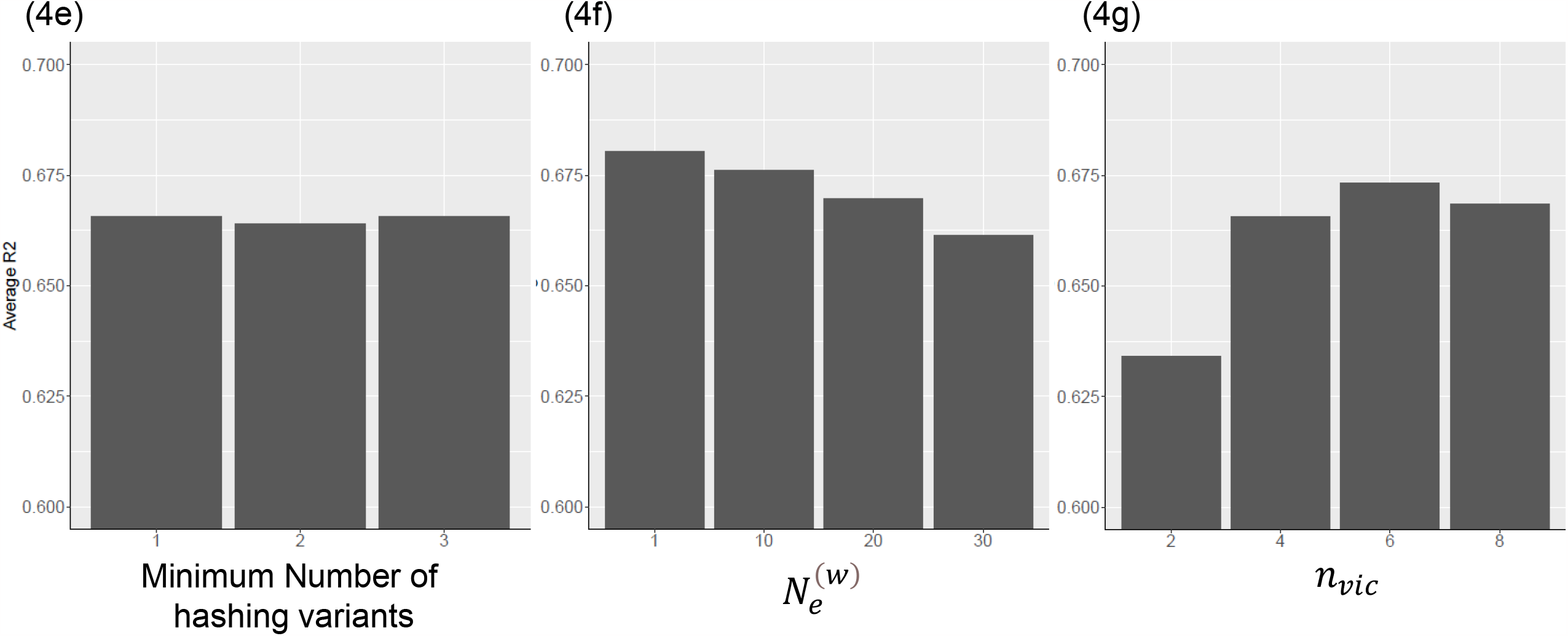
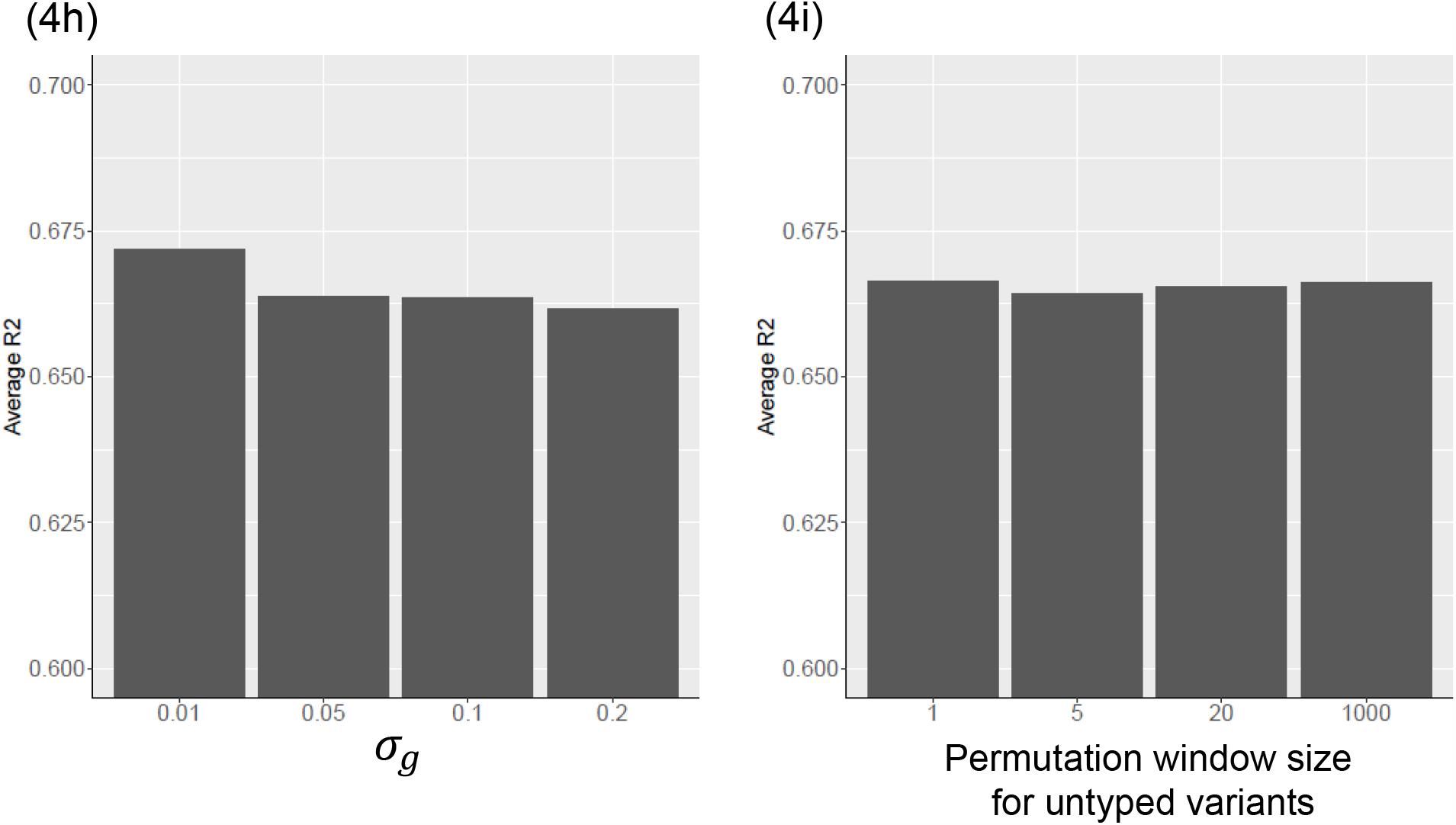
Imputation outsourcing protocol and impact of proxy generation parameters on imputation accuracy. ***(a)*** Illustration of the imputation protocol using proxy panels. Both sites use the proxy generation mechanisms and generate proxy panels. Proxy panels are sent to the outsourcing site (e.g., imputation server), which runs BEAGLE and send the imputed panel back to client site. Client generates a proxy panel from the local sensitive data (without resampling) and performs final imputation on using the results from the server. The final imputed results are mapped back to original alleles. ***(b)*** Imputation accuracy (Genotype level R2) of proxy-panel-based imputation protocol with respect to increasing 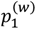, variant selection weight for first degree term in allele hashing. The imputation accuracy is measured on the African population panel. ***(c)*** Imputation accuracy vs changing 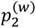, variant selection weight for 2^nd^ degree term in allele hashing. ***(d)*** Imputation accuracy vs changing 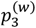, variant selection weight for 3^rd^ degree term in allele hashing. ***(e)*** Imputation accuracy vs changing minimum number of variants used in allele hashing. ***(f)*** Imputation accuracy vs increasing normalized population size, 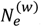, that is used in local variant selection for hashing variants. ***(g)*** Imputation accuracy vs increasing window size for allele hashing, *n*_*vic*_. ***(h)*** Imputation accuracy vs increasing standard deviation of genetic map distance noise, *σ*_*g*_. ***(i)*** Imputation accuracy vs increasing permutation of window size for untyped variants.

1. <u>Proxy-Haplotype Generation for typed variants.</u> Typed variants are used for executing the imputation models and these must be consistently encoded to generate proxy-panel for these variants. For this, the sites: (1) re-sample their panels independent of each other, (2) the sites first agree on and exchange hashing function parameters, (3) perform rolling hash-based encoding of the alleles in the panels.
2. <u>Proxy-Haplotype Generation for untyped variants.</u> Only the imputation server has access to the untyped variants. Thus, the server generates the proxy panel for untyped variants by partitioning and permutation mechanism.
3. <u>Anonymization of Variant Coordinates and Genetic Maps.</u> The server and client obfuscate the variant coordinates. Server adds noise to the genetic map after retaining only the genetic map entries for typed variants.
4. <u>Performing imputation task on Proxy Panels at 3^rd^ Party.</u> The server and client send the proxy-panels to a 3^rd^ party imputation service, which performs imputation of the variants that are untyped in client’s panel. In our scenario, we assume that this service runs BEAGLE without any modifications. We also assume that this 3^rd^ party does not have malicious intent and acts in honest-but-curious manner. These assumptions are on par with the current imputation servers and generally applicable to satisfy genomic data usage agreements.
5. <u>Sharing of the results back to Client and Re-Imputation.</u> The client downloads the imputed untyped proxy variants. It should be noted that the imputed proxy panel was originally generated using client’s re-sampled (i.e. mosaic) typed variant panel and must be processed to impute the genotypes for original panel. For this, the client encodes its original panel without the sampling step and using the rolling hash parameters that was used in the protocol. The client performs a final imputation on this proxy panel using the imputed panel from the server and obtains the untyped variant genotypes in the original panel.
6. <u>Final Decoding of Untyped Variants per Untyped Variant Sharing Policy.</u> The proxy untyped variants must be merged to build the final untyped variants. They also must be un-permuted and mapped back to their original coordinates by the client before they can be used for downstream analysis. For this, the server first decides on a policy for sharing untyped variants. This is necessary to ensure that shared variant positions do not leak any sensitive information. After deciding on sharing policy for untyped variants, the server sends the merging, allele coding, and coordinate mapping for the untyped variants. The client merges the proxy untyped variants to real untyped variants and maps them to their original positions.

### Impact of Proxy Generation Parameters on Imputation Accuracy

We first studied the impact of proxy panel generation parameters on the imputation protocol accuracy. For this test, we randomly divided the 661 subjects into two panels (330 and 331 subjects) in African population (AFR) of The 1000 Genomes Project, and extracted the 16,114 variants on chromosome 22 of Illumina Duo v3 array to be used as the typed variants. We imputed the 201,040 untyped variant genotypes on first AFR panel using the second AFR panel as the reference panel. Each parameter was varied while other parameters are held constant at default values (Methods). Imputation accuracy is estimated using genotype level R2 metric.

#### Rolling Hash Weight Selection

We first varied the parameters for random weight selection for 1^st^, 2^nd^, and 3^rd^ degree components of the rolling hash. Among these, 1^st^ degree components had the most impact on the accuracy while 2^nd^ and 3^rd^ degree component selection probabilities did not change the accuracy substantially (Fig. 4b, 4c, 4d). We chose to keep 2^nd^ and 3^rd^ degree weights to increase complexity of the hashing function. We also observed that the minimum number hash weights did not make a substantial difference in accuracy (Fig. 4e). Interestingly, we observed that increasing the variant selection weight parameter 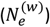, which increases probability of using more local variants for hashing each variant, decreases imputation accuracy (Fig. 4f). This result suggests that spreading the haplotype information across hashed alleles (lower variant selection parameter) increases the accuracy of imputation. The rolling hash window size (*n*_*vic*_) also has a strong impact on accuracy (Fig. 4g). We observed that increasing hashing window size increased accuracy up to 13 variants (*n*_*vic*_ = 6) and accuracy decreased at 15-variant vicinity. This suggests that spreading the allelic information on longer windows can increase accuracy. As expected, when the hashing window length becomes too large, the LD information in proxy reference panel may become spread adversely for accurate imputation.

#### Variant and Genetic Map anonymization Parameters

We assessed the two parameters that are used for anonymizing the genetic map distances (*σ*_*g*_) and untyped variant permutation filter window length. We observed that increasing *σ*_*g*_ decreases accuracy (Fig. 4h), especially between *σ*_*g*_ = 0.01 and *σ*_*g*_ = 0.05. The untyped variant permutation window size did not make a substantial difference in accuracy (Fig. 4i).This supports our expectation that HMM-based imputation tools are insensitive to the permutation of untyped variants when they are permuted between the surrounding typed variants.

### Genotype Imputation Accuracy Comparison

We first tested proxy-panel-based imputation using The 1000 Genomes Project. We randomly selected 52 subjects (2 subjects from each of the 26 populations) and extracted the genotypes as the researcher’s typed genotype panel. The typed variant positions are selected from Illumina Duo v3 array on chromosomes 19-22 (79,691 typed variants). The typed and untyped genotypes for the remaining 2,452 subjects are used as the reference panel. We executed the proxy imputation protocol with default parameters and compared the imputed genotypes with the known genotypes for the 877,387 untyped variants and calculated genotype-level R2 between known and imputed genotypes as the imputation accuracy. For comparison, we ran BEAGLE using cleartext genotypes of client’s panel and reference panel (Central Protocol), which represents the conventional outsourcing scenario with no protection of the panels. We also ran a recent method that uses resampling via Poisson point processes RESHAPE[75] to generate a synthetic imputation reference panel.

#### Comparison of Privacy Protections offered by the Methods

Central protocol serves as a baseline with no privacy protections. RESHAPE aims at protecting the reference panel and does not explicitly consider the researcher’s panel. This is by design since RESHAPE aims at increasing the distribution of reference panels with lower privacy risks. RESHAPE does not provide protection for the client’s (i.e., researcher) input panel. We also observed that the synthetic panel that RESHAPE generated perfectly preserves the allele frequency distribution of the original (sensitive) reference panel. As we have shown previously (Fig. 3c), this creates a re-identifiability concern by application of Homer’s t-statistic and LRT attacks on the synthetic panel. In retrospect, ProxyTyper’s resampling procedure does not exactly conserve the sensitive allele frequencies, although sole usage of resampling (without other mechanisms) may still cause privacy concerns (Fig. 3d). In addition, RESHAPE does not provide any mechanisms to protect the genetic maps, and variant coordinates, which renders it easier to perform the re-identification attack. Finally, the untyped variant positions in the synthetic data are clearly shared, which can lead to Bustamante’s beacon attacks with only a few variants. In comparison, ProxyTyper provides mechanisms to protect the variant coordinates, the genetic maps, and protects untyped variants by partitioning mechanism. This way, reference site can use a policy-based approach for choosing to reveal the untyped variants. We thus conclude RESHAPE and ProxyTyper to different categories in terms of the provided privacy protection wherein ProxyTyper offers a more complete and more flexible suite of protection mechanisms.

#### Comparison of Imputation Accuracy

The variants were stratified by minor allele frequency (MAF) in the whole 1000 Genomes Panel (Fig. 5a). We observed that proxy imputation protocol has lower accuracy than central protocol, the differences were minor (overall 2.7% R2, 1.8% in minor allele prediction accuracy). For example, the median accuracies were consistent among the protocols but proxy protocol predicted higher number of lower accuracy imputed variants in the rare MAF category (MAF between 0.5%-1%). For the largest MAF bin, we also observed median accuracy difference in genotype R2 was slightly higher than other MAF categories. This difference is less pronounced when we compared the minor allele prediction accuracy (Fig. 5b). Imputation using RESHAPE’s synthetic reference panel (researcher’s query panel is unprotected) provides slightly higher accuracy than proxy-based protocol (1.5% R2 and 0.08% minor allele prediction), at the cost of less privacy protection.

**Figure 5.**
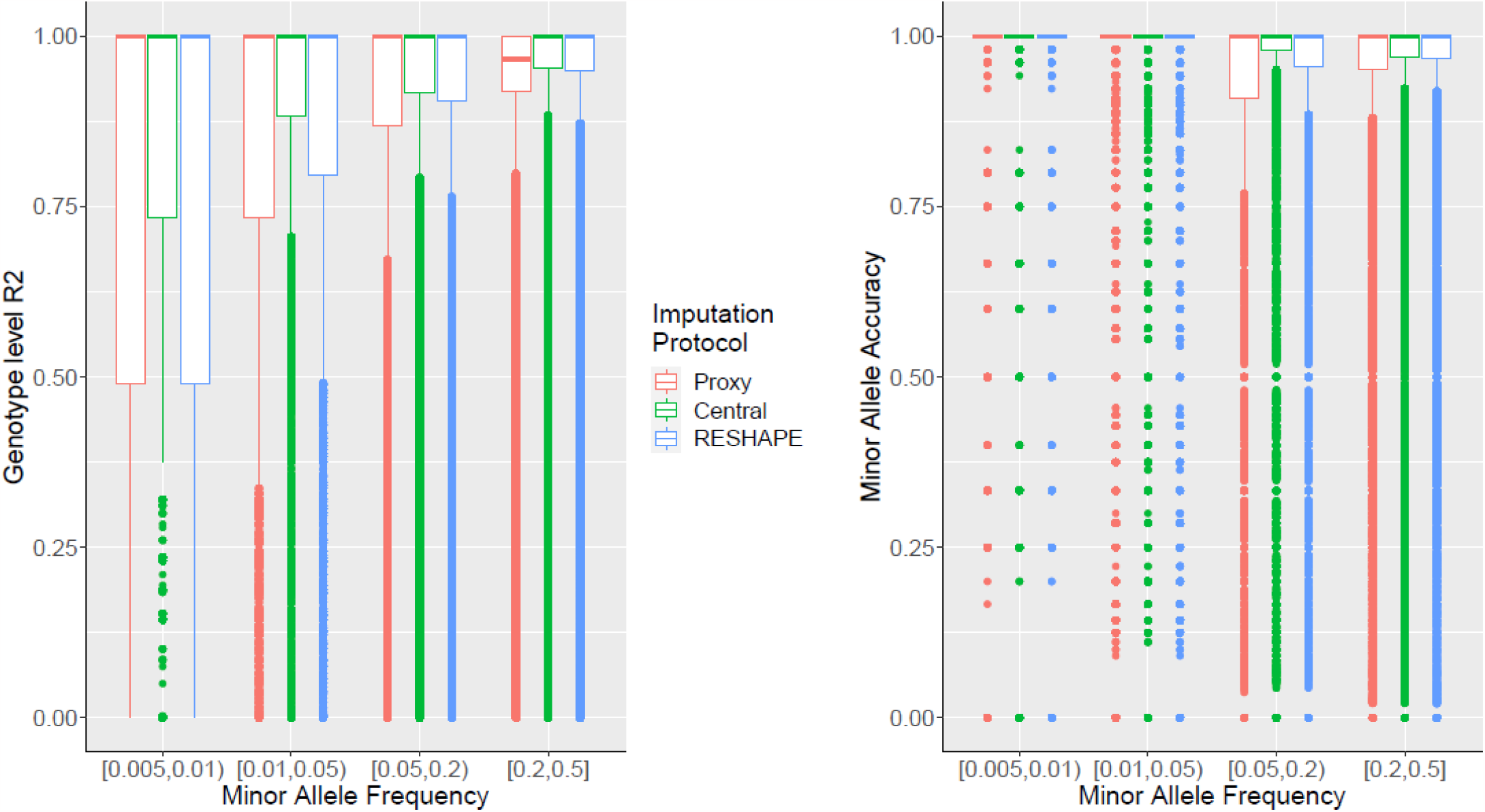
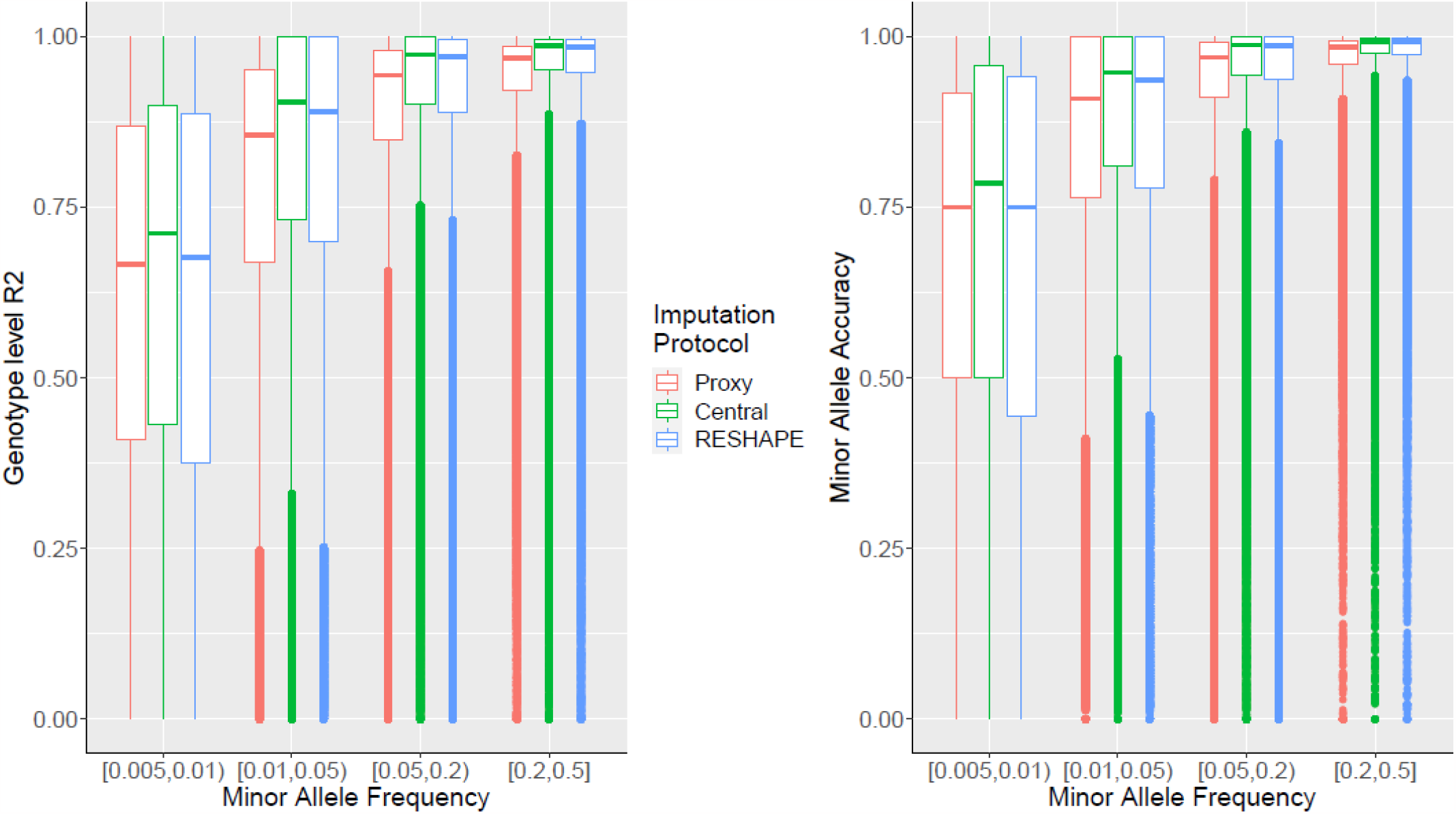
Imputation accuracy comparison between centralized, proxy, and RESHAPE. Central protocol refers to using sensitive panels in cleartext with BEAGLE. Proxy refers to the imputation protocol using proxy query (client) and reference panels as described previously. RESHAPE’s protocol uses the synthetic reference panel that is generated by RESHAPE by resampling the sensitive reference panel. With RESHAPE protocol, client’s panel is not resampled. **(a)** Comparison of the genotype level R2 (y-axis) of variants stratified with respect to different minor allele frequency (MAF) (x-axis). Imputation is performed for the randomly selected 52 subjects in 1000 Genomes Project. The remaining 2,452 subjects are used as the reference panel input to the protocols. Different methods are depicted by different colors. **(b)** Minor allele prediction accuracy of the three protocols. **(c)** Genotype R2 of imputation protocols when GTEx subjects are used as the client’s panel and 2,504 subjects in 1000 Genomes Project as the reference panel input to the protocols. **(d)** Minor allele prediction accuracy on the GTEx subjects.

We next tested the imputation with GTEx panel (635 subjects who were genotyped using whole genome sequencing) as the client set and The 1000 Genomes Project panel as the reference set. We selected the tag variants using the intersection Illumina Duo platform, GTEx variants, and the 1000 Genomes variants (54,090 variants) and the remaining variants were designated as untyped variants (450,450 variants). We again compared proxy protocol with central protocol. Similar to previous test case (52 subjects from 1000 Genomes Project), we observed that the median accuracy of central protocol is consistently higher than proxy protocol although the variance was more consistent for GTEx panel (Fig. 5c) (Overall 3.5% R2, 1.8% minor allele prediction accuracy). These differences are consistent between genotype R2 and minor allele accuracy metrics (Fig. 5d). Imputation using synthetic reference panel generated by RESHAPE exhibited slightly higher accuracy than proxy-protocol (2% in R2 and 0.07% minor allele prediction).

Overall, these results show that although proxy protocol exhibits slightly lower accuracy of imputation, it can provide a route for protecting the large-scale variant panels while utilizing existing tools with no modifications on the underlying algorithms.

## Discussion

We presented usage of proxy panels as a route to protect large scale haplotype-level datasets and demonstrated their utility for outsourcing genotype imputation. Proxy panels combine mosaic haplotype generation, noise addition, locality-sensitive random hashing, and random permutations to protect against well-known linking and re-identification of attacks. Of note, releasing rare untyped variant positions alone (even without genotype information) can lead to re-identification using Bustamante’s Beacon Attack, which are currently not considered in the literature. For high-throughput tasks such as genotype imputation, it becomes a major challenge to protect datasets against these risks. By means of proxy panel generation, the frequencies and alternate alleles of variants, variant coordinates, and genetic maps are protected with different mechanisms. These mechanisms add preemptive barriers for deterring the honest-but-curious entities (e.g., white-hat-hackers and curious researchers) and help defer accidental or intentional re-identifications through linking databases (searching of an individual in a forensic database). We therefore believe proxy-panels represent an interesting alternative route to anonymized data generation and be classified as such in context of personal data sharing regulations, e.g., GDPR and HIPAA. Another advantage of proxy panels is that they are optimized for a specific task (e.g., imputation) and are not suitable for direct secondary analyses, which comprise the basis for major privacy concerns. We are optimistic that new proxy panel generation mechanisms can be developed for optimizing the panels for task-specific scenarios that provide higher utility with low privacy concerns. One mechanism that we did not explore here is partitioning of the typed variant alleles, similar to partitioning of untyped variants. This mechanism can provide further protection against haplotype decoding by increasing the local haplotype distribution entropy beyond the mechanism that we used in this study. We foresee that new mechanisms can be designed for other tasks, e.g., kinship estimation[87], collaborative GWAS[88], as well. These mechanisms can be combined with encryption-based techniques to decrease further privacy risks while increasing efficiency.

Our results demonstrate that proxy panels can be directly used in existing imputation methods without modifications to the underlying algorithms. This is advantageous compared to, for example, homomorphic encryption-based methods, which require reformulation and careful implementation of the underlying imputation methods. In comparison, proxy panel-based methods require new protocol designs without changes in underlying algorithms, e.g., imputation task requires an extra local-imputation step at the client for obtaining the final imputation results. We believe new imputation protocol designs and post-processing tools can help increase accuracy and efficiency of proxy protocols and leave this as a future research direction.

Proxy panels should not be made publicly available since their protection rely on exchanging of secret hashing parameters between the collaborating entities (akin to symmetric encryption keys). When an adversary gains access to the hashing parameters (i.e., weights), and a proxy panel, they can use the proposed decryption strategy via haplotype-frequency analysis to decode the alleles in the proxy panel. In particular, recent studies showed that similar locality-sensitive hashing approaches (e.g., Google FLoC and Topics) may leak information when the adversary has access to the parameters of the model[89]. We foresee that proxy panels can currently enable more collaborative research (rather than public sharing of data) since they can be classified as anonymized datasets and be exchanged for analysis via non-colluding 3^rd^ parties such as AnVIL and Michigan Imputation Server. Implementing the protocols that make use of proxy panels can be seamlessly accomplished because underlying algorithms are not modified, i.e., the analysis server does not have to change their infrastructure to process proxy panel data.

Of note, the most well-known attacks in the literature are popularized because of their simplicity because they can be easily executed by honest-but-curious entities. However, these attacks require precise matching between the variants in the panels, and between the ancestral compositions of the reference panel, the target individual, and the proxy panel. Thus, proxy panels can be designed to exploit this shortcoming of the re-identification attacks: Proxy panels can be generated from randomized fractions of multi-ancestral samples. Of note, these multi-ancestral datasets can be generated once by simulations e.g., SLiM[90], or publicly available panels can be used (The 1000 Genomes Project). The tunability of proxy panel generation can be used to design new proxy-panel generation techniques that can thwart future re-identification risks with decoding attacks. Furthermore, ProxyTyper currently protects untyped variants using permutation-based anonymization. Untyped variants can be further protected using different proxy panel generation techniques wherein alleles are hashed and redundant untyped variants are excluded from computation to decrease leakage and increase efficiency. Untyped variant hashing can further make high correlations between nearby variants since untyped variants frequently share same haplotypes (i.e., linear redundancy between untyped variants) and use this correlation structure for efficient hashing. Overall, we believe there are numerous research directions that can make use of proxy-panels with minimal modifications to the underlying algorithms.

## Methods

We describe details of proxy panel generation, imputation protocol, parameter selections, and decoding attack.

### Adversarial Entities

We focus on “honest-but-curious” adversarial model[91] (most prevalent in genetic research) where users do not deviate from general protocols of analysis. In the context of collaborative genetic research, the honest-but-curious adversarial model is more prevalent and relevant than malicious entities who are actively trying to break decrypt and decode proxy panels to execute attacks. Researchers are generally assumed to be well-intentioned and follow prescribed protocols, but they might still be curious about the data they have access to. The data security protocols against malicious entities are computationally challenging to operate and maintain.

### Proxy Panel Generation

#### Mosaic Typed Variant Panel Generation by Resampling

ProxyTyper uses a re-sampling approach to generate a mosaic panel that removes the 1-to-1 correspondence of the original haplotypes to the proxy haplotype panel for providing protection against linking attacks. Given the original haplotype panel that contains *N*_*orig*_ haplotypes over *n*_*var*_ variants (denoted by *H*^(*orig*)^ matrix with *N*_*orig*_ rows and *n*_*var*_ columns), and a genetic map defined on the variant coordinates (i.e., Δ_*g*_ vector of length *n*_*var*_) as input, resampling generates *N*_*res*_ haplotypes (*H*^(*res*)^ matrix with *N*_*res*_ rows and *n*_*var*_ columns) using a Li-Stephens hidden Markov model. The sampling starts from the leftmost variant (sorted by coordinates) by selecting a random haplotype (i.e., state). At variant *i*, a new state is selected such that transition to a new state is determined by the recombination probability:

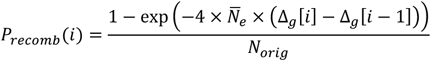

Where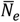 is the normalized effective population size parameter that tunes number of recombinations in the resampled panel. Probability for remaining on the same haplotype is calculated as:

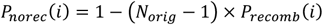

ProxyTyper samples the probabilities over the haplotypes and selects one haplotype as the new sampled haplotype. The allele on the sampled haplotype is stored as the allele for the resampled haplotype at variant *i*, i.e., 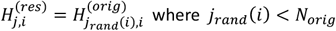 is the sampled haplotype index. After the sampling, the state is updated at variant *i* and sampling moves to the next variant. Each haplotype is sampled independently of all other haplotypes.

#### Constraints on Maximum Segment Length

For most of the variant positions no-recombination probability is larger than recombination probabilities. This may result in long consecutive allelic segments getting copied to the resampled panels. To get around this, ProxyTyper uses a parameter for constraining the length of haplotype segments copied from each haplotype. To ensure that long segments are not copied, ProxyTyper keeps track of the length of segment that is sampled from the current haplotype so far. If the length reaches a user tunable parameter 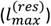, the probability of all haplotypes is set uniformly, i.e.,

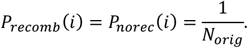

After the probabilities are reset, a new haplotype is sampled uniformly. Sampling is performed until a different haplotype is chosen similar to a rejection sampling.

#### Resampling of Untyped Variants

For the task of genotype imputation, mosaic panel generation with HMM is used for typed variants. The HMM sampling may be prohibitive for untyped variants because of their large number. To decrease the computational requirements, ProxyTyper stores the sampled haplotype indices of the typed variants at each haplotype. After haplotype *j* is generated by HMM sampling, ProxyTyper evaluates every consecutive typed variant pair. Given the typed variant pair *i* and *i* + 1, the untyped variants whose genomic coordinates are between these variants are extracted, denote by i.e., [*k, l*]. If the typed variants are sampled on the same haplotype (i.e., *j*_*rand*_(*i*) = *j*_*rand*_(*i* + 1)), the alleles for untyped variants at indices [*k, l*] are copied to the resampled haplotype (Fig. 1b, right column).

If the sampled haplotypes are different (i.e., *j*_*rand*_ (*i*) ≠ *j*_*rand*_(*i* + 1)), a recombination is introduced (Fig. 1b, left column). For this, ProxyTyper randomly selects one of the untyped variants *m* ∈ [*k, l*] with respect to their genetic positions, which represents the position of recombination. Finally, ProxyTyper copies the alleles of haplotype *j*_*rand*_(*i*) for untyped variants at indices [*k, m*] and the alleles of haplotype *j*_*rand*_(*i* + 1) at indices [*m* + 1, *l*], i.e., one transition is introduced for changing the states between typed variants *i* and *i* + 1 (Fig. 1b, right column).

#### Modulo-2 Hashing of Typed Variant Alleles

At the typed variant *i* of haplotype *j*, ProxyTyper calculates a hash of the alleles for the surrounding typed variants, 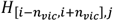 as the proxy-allele of the variant. To increase hash complexity (non-linearity), the hash includes 2^nd^ and 3^rd^ order interaction terms and a bias term:

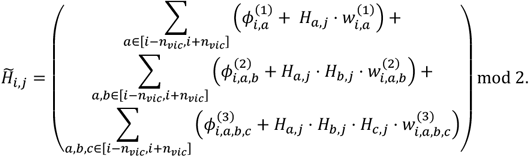

The 3 rows correspond to 1^st^, 2^nd^, and 3^rd^ degree interactions of variants at *a, b*, and *c* for variants in (2*n*_*vic*_+ 1) vicinity of variant *i*. 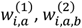, and 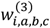 are the binary weights (specific to variant *i*) that add the corresponding component into hash. 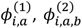, and 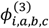 are the binary bias offsets that effectively flips the contribution of each component’s effect on the final hash. The overall hash is calculated by summing all components using modulo-2 arithmetic so that the final proxy-allele, 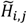, is a binary value.

#### Recombination-dependent Selection of Vicinity Variants

For variant *i*, the bias terms are selected randomly for each component using Bernoulli distribution, i.e., 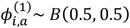. Weight parameters tune the contribution of variants in the vicinity of variant *i*, i.e., [*i* − *n*_*vic*_, *i* + *n*_*vic*_], to the proxy allele of *i*. In certain cases, the vicinity may expand a large genetic distance. To ensure that the hashes are calculated in uniform genetic vicinities, the selection of weights are performed using a genetic distance dependent manner:

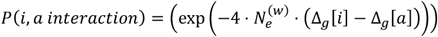

where 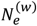 tunes the strength of interaction between pairs of variants dependent on their genetic distance, i.e., (Δ_*g*_ [*i*] − Δ_*g*_[*a*]). Low 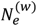 enables more relaxed selection of weights more involving variants *i* and *a* that are far from each other. This probability is used for pre-selecting the variants used for hashing the allele for variant *i*. In general, setting 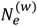 too high corresponds to focusing on immediate vicinity of variant *i*, i.e., using the variant itself for hashing.

#### Random Selection of Weights

For each weight in the hash calculation, i.e., 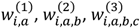, we use a Bernoulli random variable:

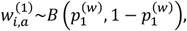

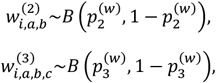

where 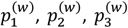 denote weight probability for 1^st^, 2^nd^, and 3^rd^ order weights, which are input arguments.

Overall, the variant selection and weight selection steps are performed independently for each variant and are parallelized. The final weights of the model include variant specific weights and biases that are stored in a binary file. This file serves as the hashing “key” and should not be shared with any entity other than the collaborating sites.

#### Untyped Variant Permutation within typed variant blocks

For the task of genotype imputation, hashing is applied to only the typed variants. This step does not protect the untyped variants of the reference panel. To conserve LD information for imputation the untyped variants between each consecutive typed variants are shuffled by their positions using a rolling permutation filter. Given the untyped variants [*k, l*] that are between the typed variants [*i, i* + 1], we permute the untyped variants indices [*k, l*] (Fig. 1d, 1e):

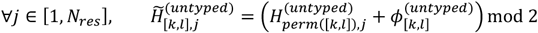

where 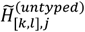 denotes the shuffled allele vector for untyped variants for variants at indices [*k, l*] on haplotype *j*, and *perm* ([*k, l*]) denotes a random permutation of indices *k, k* + 1, …, *l* − 1, 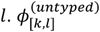 denotes a random binary bias that randomly flips the alternate alleles of untyped variants in [*k, l*]. Each entry in the bias vector is selected with a Bernoulli variable, i.e., *B*(0.5,0,5). To tune the extent of index permutation, ProxyTyper performs permutations recursively on sliding windows of length 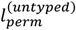over the set of untyped variants [*k, l*]. Smaller 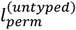 better preserves the ordering of the untyped variants.

#### Generation of Proxy Untyped Variants by Allele Partitioning

Although permutation obfuscates variant positions, it precisely preserves the allele frequencies of the untyped variants. ProxyTyper generate proxy untyped variants by a mechanism we termed “Allele Partitioning”. Given an untyped variant, the basic idea of partitioning is randomly splitting the alleles among two new proxy variants such that the union of the proxy variants exactly recapitulates the alleles in the original variant (Fig. 1d):

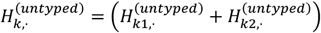

where we treat the alleles of *k*^*th*^ untyped variant as a binary vector across haplotypes. This vector is equal to the summation of two proxy untyped variant allele vectors, 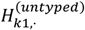 and 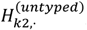, which denote the (binary) allele vectors of the proxy untyped variants *k*1 and *k*2. The above equation simply refers to partitioning the haplotypes that harbor alternative alleles for untyped variant *k*. For each original untyped variant in the reference panel, ProxyTyper generates two untyped proxy variants by randomly partitioning the haplotypes that harbor alternate allele for the untyped variant. The original untyped variant is included in the proxy untyped reference panel anymore and it is replaced with two proxy untyped variants, which are randomly placed between the nearest tag variants on the left and right. After partitioning, the proxy untyped alleles always have smaller allele frequencies than the original variant.

The motivation for using this partitioning as a mechanism is that it obfuscates the alleles of the original untyped variant *k* by partitioning its alleles to two proxy variants but it preserves the imputed allelic probabilities in imputation tasks. For example, given *k*^*th*^ untyped variant, the imputed probability of allele 1 is equal to the summation of the HMM probabilities of all haplotypes that harbor allele 1 on this variant:

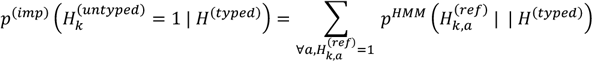

*p*^(*imp*)^ denotes the imputed alternate allele probability of *k*^*th*^ variant by BEAGLE and 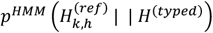 denotes the imputation HMM’s forward-backward state probability for haplotype *h* at variant *k*, where *h*^*th*^ haplotype of the reference panel harbors an alternate allele for variant *k*.

When we partition *k*^*th*^ variant’s alleles into two proxy untyped variants (k1 and k2), we effectively partition the haplotypes that harbor an alternate allele for variant *k* to these two variants:

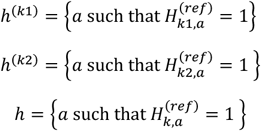

and

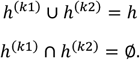

The last equations hold by the partitioning of the alleles such that 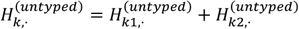. The partitioning of haplotypes allows us to reconstruct the imputed alternate allele probability for variant *k* in terms of the alternate allele probabilities assigned to *k*1 and *k*2. We first decompose the haplotypes in the equation for alternate allele probability at variant *k*:

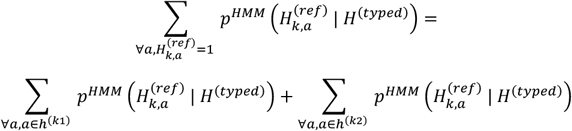

Given the panel with proxy untyped variants *k*1 and *k*2, we can approximate the HMM haplotype state probabilities using the imputation performed with panel proxy untyped:

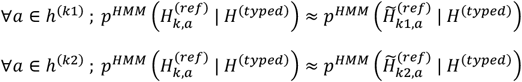

These equations hold approximately because the state probability at haplotype *a* of imputation HMM at untyped variants rely only on the closest typed variants on the left and right vicinity of the untyped variant. This is satisfied because *k*1 and *k*2 are constrained to be between the same typed variants as *k* (Fig. 1d).

Left hand side of the equations refer to the BEAGLE’s HMM probability for the haplotype *a* when we perform imputation using typed variants as input and the original reference panel. Right hand side indicates the same probability when we use the untyped reference panel with proxy variants *k*1 and *k*2, which are new untyped variants that do not exist in the original panel. Replacing the right-hand side of the equations with the above we get:

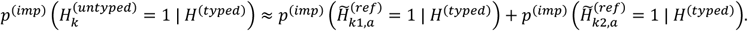

This equation indicates that we can use the reference panel with proxy untyped variants 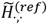, obtain the imputed alternate allele probabilities (using BEAGLE) for proxy untyped variants at *k*1 and *k*2, and finally map them back to the original untyped variant *k* by summing their probabilities. ProxyTyper uses one more mechanism to further obfuscate the proxy untyped variants by randomly flipping their alleles with 50% probability. This operation does not accrue any accuracy penalty since it also effectively flips the probability of alternate alleles. Furthermore, allele flipping helps obfuscate the possible linkage between original and proxy untyped variants because the proxy untyped variants can have higher frequency than that original variant. If a proxy untyped variant is flipped, ProxyTyper takes this into consideration by subtracting it from one before mapping it back to original untyped variant.

Currently, ProxyTyper can partition each untyped variants into 2 proxy untyped variants, which doubles the number of untyped variants in the proxy panel. Each proxy untyped variant is independently flipped. The partitioning information is efficiently stored in one file (kept secret at reference site) that describe which proxy variants correspond to which original variants and the flipping states of each proxy variant. Note that each untyped proxy variant is placed to a random position between the surrounding tag variants, similar to the permutation mechanism.

### Anonymization of Genetic Maps and Variant Coordinates

After sampling of haplotypes and hashing of alleles, ProxyTyper first maps variant coordinates (including typed and untyped variants) to a uniform range. This is set simply to 100 megabases be default. It should be noted that the variant coordinates are only used for sorting variants and do not change imputation accuracy since genetic maps are provided. After coordinate anonymization, the genetic maps are censored to release the genetic distances for only the typed variants. For this, ProxyTyper first extracts the original cumulative genetic distances for only the typed variants. Next, a Gaussian noise is added to the genetic distances:

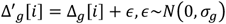

where, *σ*_*g*_ is a user-defined standard deviation of genetic distance noise. Next, 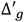 are sorted over all typed variants and finally assigned as the anonymized coordinates:

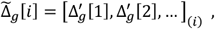

where 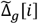 denotes the anonymized genetic distance for typed variant *i* and 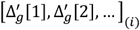 denotes the *i*^*th*^ element in the sorted sequence of noisy genetic distances.

### Viterbi Decoding of Attack on Typed Variants and Likelihood Ratio Test Reidentification

We describe the Viterbi decoding algorithm that is used for decoding the hashed alleles in the proxy panels. Decoding algorithm is similar to an imputation HMM that decodes the hashed (proxy) alleles in a proxy panel by combining a frequency analysis of proxy panel k-mers with reference panel k-mers. It should be noted that the k-mer length in decoding is independent of the proxy panel generation k-mer lengths.

### Inputs

The adversary has access to a proxy panel that is being decoded. Adversary uses a public reference panel to estimate the frequencies of the cleartext k-mers and match them at each k-mer. It should also be noted that the proxy panel’s variant coordinates are anonymized, i.e., adversary needs to predict the positions of the typed variants. Rather than predicting the positions of variants, we assumed that the attacker approximately predicts where the variants are using, for example, LD statistics. To simulate this, we divided the 87,960 typed variants into 2 equal sized sets by assigning every 1^st^ variant to the first set and every 2^nd^ variant to the second set, i.e., the variants sets are interleaved with respect to each other. Coordinates of the 1^st^ variant set are used for the proxy panel and those of the 2^nd^ set is used for the reference panel. Simply put, this corresponds to the attacker using an interleaved set of variants for the attack. The proxy panel is generated from a sample of 100 subjects from the AFR populations in the 1000 Genomes Project. For decreasing computational complexity, we focused on a region of 100 megabases (chr1:10,000,000-110,000,000), which contains 19,379 variants. The reference panel (in comprises the genotypes of 461 subjects. We also used a hold-out panel of 100 subjects that are used as a control panel.

### Transition and Emission Probabilities

The decoding HMM uses a modification of the Li-Stephens model[84] that is used in imputation methods. The decoding HMM runs on the coordinates of the reference panel and uses each reference haplotype as a state in the Markov model. The transition probability at each position is assigned using the genetic distances and the consecutive k-mer concordance. At reference variant position *i*, the transition probability is defined as:

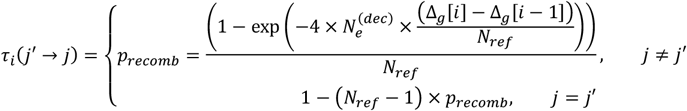

which is described by Li-Stephens model. Note that the transition does not rely on the haplotype indices (i.e., states) except for the staying on the same state, i.e., all same state transitions are assigned the same probability. 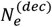 is set to default value of 10^6^. We modify this transition matrix to introduce the concordance of allele frequencies:

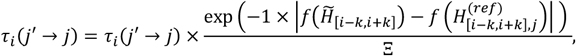

where *k* denotes the vicinity length (i.e., total k-mer length is (2*k* + 1)) and Ξ denotes the normalization factor over all haplotypes. Above equation integrates concordance of the allele frequencies between the proxy panel and the reference panel to the transition score. We further need to introduce the constraint on consistency of the consecutive k-mers on the current haplotype (i.e., state *j*) state and previous state (state *k*) at variant position *i* − 1:

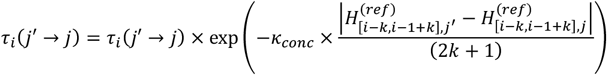

where we score the distance between the overlapping substrings of the two k-mers at positions [*i* − 1 − *k, i* − 1 + *k*] and [*i* − *k, i* + *k*]. The modifications in the equations constrain the concordance between the frequencies of the reference panel and the proxy panel, and also the concordance of the emitted reference panel k-mers between current haplotype and the previous haplotype. *κ*_*conc*_ denotes a weight that tunes the weight of k-mer concordance between two positions. This value is set as:

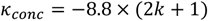

which penalizes every additional mismatch between the consecutive k-mers by 10^−4^, which is the default mismatch score in BEAGLE for allelic errors.

### Scoring Matrix Calculation

Given a reference panel haplotype that consists of *n*_*var*_ variants, we allocate *n*_*var*_ × *N*_*ref*_ matrix of probability scores, denoted by *S*. At variant position *i*, the score at state (i.e., haplotype) *j* is calculated by evaluating all state transitions at the previous variant position *i* − 1:

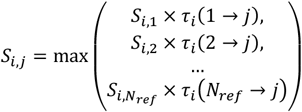

This recursion is calculated for all reference haplotypes *j* < *N*_*ref*_. The scoring matrix is calculated for each proxy haplotype over all variants. The final score matrix is traced back using Viterbi algorithm to identify the highest scoring haplotype sequence over the reference haplotypes and is used as the decoded allele sequence.

### Reidentification of individuals within decoded proxy panel using LRT Attack

After the alleles are decoded, we use the decoded haplotype matrix as a pool in which a target individual with known genome is searched. The adversary uses the same reference panel used for performing the re-identification attack. The LRT attack statistic is calculated as described in Sankararaman et. al. (Supplementary Information page 25 in [39]) by comparing the allele frequencies of variants between the pool and the reference dataset. LRT statistic is calculated as:

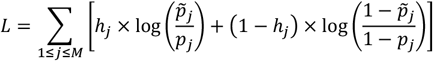

where *h*_*j*_ ∈ {0,1} denotes the binary allele value at *j*^*th*^ variant, 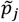 denotes the frequency of alternate allele (*h*_*j*_ = 1) in the pool and *p*_*j*_ denotes the frequency of alternate allele in the reference panel. LRT statistics are calculated for all decoded haplotypes.

We evaluated three case scenarios to calculate the re-identification statistics that are shown in Figure 3b, 3c, 3d:

1. Use decoded proxy panel as the pool (Actual attack scenario, Fig. 3b): This is the basic scenario when proxy panel is used for protecting the panel. Adversary tries to identify if an individual with known genotypes is in the decoded panel.
2. Use the cleartext original panel as the pool (Baseline scenario with original data for pool panel with no protection, Fig. 3c): This is the baseline attack scenario where adversary has access to the cleartext sensitive panel and tries to identify individual’s participation in the panel.
3. Use the cleartext resampled original panel as the pool (Control case scenario with sampled mosaic panel without allele hashing as the pool panel, Fig. 3d): This scenario evaluates the participation prediction when only resampling is used.

## REFERENCES

1. Muir P, Li S, Lou S, Wang D, Spakowicz DJ, Salichos L, et al. The real cost of sequencing: scaling computation to keep pace with data generation. Genome Biol. 2016;17: 53.

2. Chen W, Coombes BJ, Larson NB. Recent advances and challenges of rare variant association analysis in the biobank sequencing era. Front Genet. 2022;13: 1014947.

3. Locke DP, Sharp AJ, McCarroll SA, McGrath SD, Newman TL, Cheng Z, et al. Linkage disequilibrium and heritability of copy-number polymorphisms within duplicated regions of the human genome. Am J Hum Genet. 2006;79: 275–290.

4. International HapMap Consortium. A haplotype map of the human genome. Nature. 2005;437: 1299–1320.

5. The 1000 Genomes Project Consortium. A global reference for human genetic variation. Nature. 2015. pp. 68–74. doi:10.1038/nature15393

6. Kowalski MH, Qian H, Hou Z, Rosen JD, Tapia AL, Shan Y, et al. Use of >100,000 NHLBI Trans-Omics for Precision Medicine (TOPMed) Consortium whole genome sequences improves imputation quality and detection of rare variant associations in admixed African and Hispanic/Latino populations. PLoS Genet. 2019;15. doi:10.1371/journal.pgen.1008500

7. All of Us Research Program Investigators, Denny JC, Rutter JL, Goldstein DB, Philippakis A, Smoller JW, et al. The “All of Us” Research Program. N Engl J Med. 2019;381: 668–676.

8. Popejoy AB, Fullerton SM. Genomics is failing on diversity. Nature. Nature Publishing Group; 2016. pp. 161–164. doi:10.1038/538161a

9. Matalon DR, Zepeda-Mendoza CJ, Aarabi M, Brown K, Fullerton SM, Kaur S, et al. Clinical, technical, and environmental biases influencing equitable access to clinical genetics/genomics testing: A points to consider statement of the American College of Medical Genetics and Genomics (ACMG). Genet Med. 2023; 100812.

10. Bentley AR, Callier S, Rotimi CN. Diversity and inclusion in genomic research: why the uneven progress? J Community Genet. 2017;8: 255–266.

11. Choudhury A, Aron S, Botigué LR, Sengupta D, Botha G, Bensellak T, et al. High-depth African genomes inform human migration and health. Nature. 2020;586: 741–748.

12. Stark Z, Boughtwood T, Phillips P, Christodoulou J, Hansen DP, Braithwaite J, et al. Australian Genomics: A federated model for integrating genomics into healthcare. Am J Hum Genet. 2019;105: 7–14.

13. Bonomi L, Huang Y, Ohno-Machado L. Privacy challenges and research opportunities for genomic data sharing. Nat Genet. 2020;52: 646–654.

14. Yousefi S, Abbassi-Daloii T, Kraaijenbrink T, Vermaat M, Mei H, van’t Hof P, et al. A SNP panel for identification of DNA and RNA specimens. BMC Genomics. 2018;19. doi:10.1186/s12864-018-4482-7

15. Hafiza MK, Sajjad A, Nasir S, Qazi LA, Muhammad A, Mohammad AT. Exoneration of primary suspect after false confession with the help of forensic DNA analysis. Forensic Genom. 2022;2: 17–20.

16. Seydel C. You can run, but your DNA can’t hide. Forensic Genom. 2022;2: 97–102.

17. Niemiec E, Howard HC. Ethical issues in consumer genome sequencing: Use of consumers’ samples and data. Appl Transl Genom. 2016;8: 23–30.

18. Pulivarti R. Cybersecurity of Genomic Data. Gaithersburg, MD: National Institute of Standards and Technology; 2023. doi:10.6028/nist.ir.8432.ipd

19. Sherburn IA, Finlay K, Best S. How does the genomic naive public perceive whole genomic testing for health purposes? A scoping review. Eur J Hum Genet. 2023;31: 35–47.

20. Jamal L, Sapp JC, Lewis K, Yanes T, Facio FM, Biesecker LG, et al. Research participants’ attitudes towards the confidentiality of genomic sequence information. Eur J Hum Genet. 2014;22: 964–968.

21. After Havasupai litigation, Native Americans wary of genetic research. Am J Med Genet A. 2010;152A: fmix.

22. Garrison NA. Genomic justice for native Americans: Impact of the Havasupai case on genetic research. Sci Technol Human Values. 2013;38: 201–223.

23. Kang JTL, Goldberg A, Edge MD, Behar DM, Rosenberg NA. Consanguinity rates predict long runs of homozygosity in Jewish populations. Hum Hered. 2016;82: 87–102.

24. Powell K. The broken promise that undermines human genome research. Nature. 2021;590: 198–201.

25. Walking the tightrope between data sharing and data protection. Nat Med. 2022;28: 873.

26. Budin-Ljøsne I, Isaeva J, Knoppers BM, Tassé AM, Shen H-Y, McCarthy MI, et al. Data sharing in large research consortia: experiences and recommendations from ENGAGE. Eur J Hum Genet. 2014;22: 317–321.

27. Stobbe MD, Gonzalez-Perez A, Lopez-Bigas N, Gut IG. Ten simple rules for a successful international consortium in big data omics. PLoS Comput Biol. 2022;18: e1010546.

28. General Data Protection Regulation (GDPR) Compliance Guidelines. In: GDPR.eu [Internet]. [cited 5 May 2020]. Available: https://gdpr.eu/

29. Cohen IG, Mello MM. HIPAA and Protecting Health Information in the 21st Century. JAMA. 2018;320: 231–232.

30. Greenbaum D, Sboner A, Mu XJ, Gerstein M. Genomics and privacy: Implications of the new reality of closed data for the field. PLoS Computational Biology. 2011. doi:10.1371/journal.pcbi.1002278

31. Hubaux J-P, Katzenbeisser S, Malin B. Genomic data privacy and security: Where we stand and where we are heading. IEEE Secur Priv. 2017;15: 10–12.

32. Wan Z, Hazel JW, Clayton EW, Vorobeychik Y, Kantarcioglu M, Malin BA. Sociotechnical safeguards for genomic data privacy. Nat Rev Genet. 2022;23: 429–445.

33. Erlich Y, Narayanan A. Routes for breaching and protecting genetic privacy. Nat Rev Genet. 2014;15: 409–421.

34. Erlich Y, Williams JB, Glazer D, Yocum K, Farahany N, Olson M, et al. Redefining genomic privacy: trust and empowerment. PLoS Biol. 2014;12: e1001983.

35. Shabani M, Marelli L. Re-identifiability of genomic data and the GDPR: Assessing the re-identifiability of genomic data in light of the EU General Data Protection Regulation. EMBO Rep. 2019;20: e48316.

36. Gymrek M, McGuire AL, Golan D, Halperin E, Erlich Y. Identifying personal genomes by surname inference. Science. 2013;339: 321–324.

37. Lin Z, Owen AB, Altman RB. Genetics. Genomic research and human subject privacy. Science. 2004;305: 183.

38. Homer N, Szelinger S, Redman M, Duggan D, Tembe W, Muehling J, et al. Resolving individuals contributing trace amounts of DNA to highly complex mixtures using high-density SNP genotyping microarrays. PLoS Genet. 2008;4: e1000167.

39. Sankararaman S, Obozinski G, Jordan MI, Halperin E. Genomic privacy and limits of individual detection in a pool. Nat Genet. 2009;41: 965–967.

40. Visscher PM, Hill WG. The limits of individual identification from sample allele frequencies: theory and statistical analysis. PLoS Genet. 2009;5: e1000628.

41. Im HK, Gamazon ER, Nicolae DL, Cox NJ. On sharing quantitative trait GWAS results in an era of multiple-omics data and the limits of genomic privacy. Am J Hum Genet. 2012;90: 591–598.

42. Harmanci A, Gerstein M. Quantification of private information leakage from phenotype-genotype data: linking attacks. Nat Methods. 2016;13: 251–256.

43. Harmanci A, Gerstein M. Analysis of sensitive information leakage in functional genomics signal profiles through genomic deletions. Nat Commun. 2018;9. doi:10.1038/s41467-018-04875-5

44. Branum R, Wolf SM. International policies on sharing genomic research results with relatives: Approaches to balancing privacy with access. J Law Med Ethics. 2015;43: 576–593.

45. Telenti A, Ayday E, Hubaux JP. On genomics, kin, and privacy. F1000Res. 2014. doi:10.12688/f1000research.3817.1

46. Bu D, Wang X, Tang H. Haplotype-based membership inference from summary genomic data. Bioinformatics. 2021;37: i161–i168.

47. Jacobs KB, Yeager M, Wacholder S, Craig D, Kraft P, Hunter DJ, et al. A new statistic and its power to infer membership in a genome-wide association study using genotype frequencies. Nat Genet. 2009;41: 1253–1257.

48. Fiume M, Cupak M, Keenan S, Rambla J, de la Torre S, Dyke SOM, et al. Federated discovery and sharing of genomic data using Beacons. Nat Biotechnol. 2019;37: 220–224.

49. Shringarpure SS, Bustamante CD. Privacy Risks from Genomic Data-Sharing Beacons. Am J Hum Genet. 2015;97: 631–646.

50. Ayoz K, Ayday E, Cicek AE. Genome reconstruction attacks against genomic data-sharing beacons. Proc Priv Enhancing Technol. 2021;2021: 28–48.

51. Thenen NV, Ayday E, Cicek AE. Re-Identification of Individuals in Genomic Data-Sharing Beacons via Allele Inference. Bioinformatics. 2018. doi:10.1101/200147

52. Egeland T, Fonneløp AE, Berg PR, Kent M, Lien S. Complex mixtures: a critical examination of a paper by Homer et al. Forensic Sci Int Genet. 2012;6: 64–69.

53. Sampson J, Zhao H. Identifying individuals in a complex mixture of DNA with unknown ancestry. Stat Appl Genet Mol Biol. 2009;8: Article 37.

54. Dwork C, Roth A. The algorithmic foundations of differential privacy. Found Trends Theor Comput Sci. 2013;9: 211–407.

55. Dwork C. Differential privacy: A cryptographic approach to private data analysis. Privacy, Big Data, and the Public Good. New York: Cambridge University Press; 2014. pp. 296–322.

56. Kim M, Harmanci AO, Bossuat J-P, Carpov S, Cheon JH, Chillotti I, et al. Ultrafast homomorphic encryption models enable secure outsourcing of genotype imputation. Cell Systems. 2021;12: 1108–1120.e4.

57. Yang M, Zhang C, Wang X, Liu X, Li S, Huang J, et al. TrustGWAS: A full-process workflow for encrypted GWAS using multi-key homomorphic encryption and pseudorandom number perturbation. Cell Syst. 2022;13: 752–767.e6.

58. Froelicher D, Troncoso-Pastoriza JR, Raisaro JL, Cuendet MA, Sousa JS, Cho H, et al. Truly privacy-preserving federated analytics for precision medicine with multiparty homomorphic encryption. Nat Commun. 2021;12: 5910.

59. TrustGWAS: A full-process workflow for encrypted genome-wide association studies using multi-key homomorphic encryption and pseudo-random number perturbation. Github; Available: https://github.com/melobio/TrustGWAS

60. Cho H, Wu DJ, Berger B. Secure genome-wide association analysis using multiparty computation. Nat Biotechnol. 2018;36: 547–551.

61. Blatt M, Gusev A, Polyakov Y, Rohloff K, Vaikuntanathan V. Optimized homomorphic encryption solution for secure genome-wide association studies. BMC Med Genomics. 2020;13: 83.

62. Shimizu K, Nuida K, Rätsch G. Efficient privacy-preserving string search and an application in genomics. Bioinformatics. 2016;32: 1652–1661.

63. Nakagawa Y, Ohata S, Shimizu K. Efficient privacy-preserving variable-length substring match for genome sequence. Algorithms Mol Biol. 2022;17: 9.

64. Popic V, Batzoglou S. A hybrid cloud read aligner based on MinHash and kmer voting that preserves privacy. Nat Commun. 2017. doi:10.1038/ncomms15311

65. Gentry C. A FULLY HOMOMORPHIC ENCRYPTION SCHEME. PhD Thesis. 2009; 1–209.

66. Kim M, Lauter K. Private genome analysis through homomorphic encryption. BMC Med Inform Decis Mak. 2015;15 Suppl 5: S3.

67. Dowlin N, Gilad-Bachrach R, Laine K, Lauter K, Naehrig M, Wernsing J. Manual for using homomorphic encryption for bioinformatics. Proc IEEE Inst Electr Electron Eng. 2017;105: 1–16;

68. Orlandi C. Is multiparty computation any good in practice? ICASSP, IEEE International Conference on Acoustics, Speech and Signal Processing - Proceedings. 2011. doi:10.1109/ICASSP.2011.5947691

69. Zhao C, Zhao S, Zhao M, Chen Z, Gao C-Z, Li H, et al. Secure Multi-Party Computation: Theory, practice and applications. Inf Sci (Ny). 2019;476: 357–372.

70. Wang W, Chen G, Pan X, Zhang Y, Wang XF, Bindschaedler V, et al. Leaky cauldron on the dark land: Understanding memory side-channel hazards in SGX. Proceedings of the ACM Conference on Computer and Communications Security. New York, NY, USA: Association for Computing Machinery; 2017. pp. 2421–2434.

71. Nilsson A, Bideh PN, Brorsson J. A survey of published attacks on Intel SGX. arXiv. 2020. Available: http://arxiv.org/abs/2006.13598

72. Bernier A, Liu H, Knoppers BM. Computational tools for genomic data de-identification: facilitating data protection law compliance. Nat Commun. 2021;12: 6949.

73. Heeney C, Hawkins N, de Vries J, Boddington P, Kaye J. Assessing the privacy risks of data sharing in genomics. Public Health Genomics. 2011;14: 17–25.

74. Gonzales A, Guruswamy G, Smith SR. Synthetic data in health care: A narrative review. PLOS Digit Health. 2023;2: e0000082.

75. Cavinato T, Rubinacci S, Malaspinas A-S, Delaneau O. A resampling-based approach to share reference panels. bioRxiv. 2023. p. 2023.04.07.535812. doi:10.1101/2023.04.07.535812

76. Yelmen B, Decelle A, Ongaro L, Marnetto D, Tallec C, Montinaro F, et al. Creating artificial human genomes using generative neural networks. PLoS Genet. 2021;17: e1009303.

77. Wohns AW, Wong Y, Jeffery B, Akbari A, Mallick S, Pinhasi R, et al. A unified genealogy of modern and ancient genomes. Science. 2022;375: eabi8264.

78. Anderson-Trocmé L, Nelson D, Zabad S, Diaz-Papkovich A, Kryukov I, Baya N, et al. On the genes, genealogies, and geographies of Quebec. Science. 2023;380: 849–855.

79. Sun Q, Liu W, Rosen JD, Huang L, Pace RG, Dang H, et al. Leveraging TOPMed imputation server and constructing a cohort-specific imputation reference panel to enhance genotype imputation among cystic fibrosis patients. HGG Adv. 2022;3: 100090.

80. Magrabi F, Ong MS, Coiera E. Health IT for patient safety and improving the safety of health IT. Stud Health Technol Inform. 2016;222. Available: https://pubmed.ncbi.nlm.nih.gov/27198089/

81. Song M, Greenbaum J, Luttrell J IV, Zhou W, Wu C, Luo Z, et al. An autoencoder-based deep learning method for genotype imputation. Front Artif Intell. 2022;5. doi:10.3389/frai.2022.1028978

82. Dias R, Evans D, Chen S-F, Chen K-Y, Loguercio S, Chan L, et al. Rapid, Reference-Free human genotype imputation with denoising autoencoders. Elife. 2022;11: e75600.

83. Yu K, Das S, LeFaive J, Kwong A, Pleiness J, Forer L, et al. Meta-imputation: An efficient method to combine genotype data after imputation with multiple reference panels. Am J Hum Genet. 2022;109: 1007–1015.

84. Li N, Stephens M. Modeling Linkage Disequilibrium and Identifying Recombination Hotspots Using Single-Nucleotide Polymorphism Data. Genetics. 2003;165: 2213–2233.

85. Ycart B. Letter counting: a stem cell for Cryptology, Quantitative Linguistics, and Statistics. arXiv [math.HO]. 2012. Available: http://arxiv.org/abs/1211.6847

86. Wang S, Kim M, Jiang X, Harmanci AO. Evaluation of vicinity-based hidden Markov models for genotype imputation. BMC Bioinformatics. 2022;23. doi:10.1186/s12859-022-04896-4

87. Wang S, Kim M, Li W, Jiang X, Chen H, Harmanci A. Privacy-aware estimation of relatedness in admixed populations. Brief Bioinform. 2022;23. doi:10.1093/bib/bbac473

88. Li W, Chen H, Jiang X, Harmanci A. Federated generalized linear mixed models for collaborative genome-wide association studies. iScience. 2023;26: 107227.

89. Turati F, Cotrini C, Kubicek K, Basin D. Locality-sensitive hashing does not guarantee privacy! Attacks on Google’s FLoC and the MinHash Hierarchy system. arXiv [cs.CR]. 2023. Available: http://arxiv.org/abs/2302.13635

90. Haller BC, Messer PW. SLiM 3: Forward genetic simulations beyond the Wright-Fisher model. Mol Biol Evol. 2019;36: 632–637.

91. Paverd A, Martin A, Brown I. Modelling and automatically analysing privacy properties for honest-but-curious adversaries. [cited 31 May 2023]. Available: https://www.cs.ox.ac.uk/people/andrew.paverd/casper/casper-privacy-report.pdf

